# Transposon-sequencing across multiple *Mycobacterium abscessus* isolates reveals significant functional genomic diversity among strains

**DOI:** 10.1101/2023.03.29.534805

**Authors:** Chidiebere Akusobi, Sanjeevani Choudhery, Bouchra S. Benghomari, Ian D. Wolf, Shreya Singhvi, Thomas R. Ioerger, Eric J. Rubin

## Abstract

*Mycobacterium abscessus (Mab)* is a clinically significant pathogen and a highly genetically diverse species due to its large accessory genome. The functional consequence of this diversity remains unknown mainly because, to date, functional genomic studies in *Mab* have been primarily performed on reference strains. Given the growing public health threat of *Mab* infections, understanding the functional genomic differences among *Mab* clinical isolates can provide more insight into how its genetic diversity influences gene essentiality, clinically relevant phenotypes, and importantly, potential drug targets. To determine the functional genomic diversity among *Mab* strains, we conducted transposon-sequencing (TnSeq) on 21 genetically diverse clinical isolates, including 15 *M. abscessus subsp. abscessus* isolates and 6 *M. abscessus subsp. massiliense* isolates, cataloging all the essential and non-essential genes in each strain. Pan-genome analysis revealed a core set of 3845 genes and a large accessory genome of 11,507. We identified 259 core essential genes across the 21 clinical isolates and 425 differentially required genes, representing ∼10% of the *Mab* core genome. We also identified genes whose requirements were sub-species, lineage, and isolate-specific. Finally, by correlating TnSeq profiles, we identified 19 previously uncharacterized genetic networks in *Mab*. Altogether, we find that *Mab* clinical isolates are not only genetically diverse but functionally diverse as well.

## Introduction

*Mycobacterium abscessus (Mab)* has emerged in recent years as a growing threat to public health due to the rise of *Mab* infections worldwide (1). *Mab* causes a wide range of infections including 80% of pulmonary infections from rapidly growing mycobacteria (2,3). Treating *Mab* infections is difficult due to *Mab’s* high level of intrinsic resistance to many antibiotics, including most of the cornerstones of tuberculosis treatment (4–7). Importantly, treating *Mab* requires identifying the subspecies of the clinical isolate, which influences antibiotic treatment based on the presence of *erm*, a gene that confers inducible macrolide resistance (8,9).

*Mab* is divided into three subspecies: *abscessus, bolletii,* and *massiliense* (10,11). Notably, most *massiliense* isolates lack an active *erm* gene (8,9). Whole-genome sequencing of *Mab* clinical isolates revealed that *Mab* has a large, open pan-genome, with isolates sharing ∼75% of their genome in what is known as the ‘core genome’ (12,13). The remaining 25% of gene content is in the ‘accessory genome’ defined as genes present in some clinical isolates but not all. Most accessory genome content comprises plasmids, prophages, and genomic islands acquired by horizontal gene transfer (HGT) (14). Unlike *Mycobacterium tuberculosis (Mtb),* which evolves clonally likely due to its facultative intracellular lifestyle (15), the primary driver of *Mab* evolution is recombination and HGT (12,16,17). As a result, compared to TB, *Mab* clinical isolates have larger accessory genomes, which are significant contributors to the genetic diversity present in this species.

While *Mab* clinical isolates have large accessory genomes, the functional consequence of this diversity remains unknown. Previous studies have shown that accessory gene content impacts genetic networks, generating new phenotypes, gene dependencies, and redundancies within strains (18–20). Variation in accessory genomes likely generates functional genomic and phenotypic differences between isolates. The functional differences between isolates can be understood through genetic screens using CRISPRi and transposon-sequencing (TnSeq) libraries. TnSeq and the subsequent identification of essential genes have typically been conducted in lab-adapted reference strains. However, in recent years, groups have performed TnSeq on clinical isolates of pathogens, including *S. enterica, S. pneumoniae, S. aureus,* and *M. tuberculosis.* These studies identified genes that were differentially essential among clinical isolates and that these differences influenced clinically relevant phenotypes (19, 21–24).

For *Mab*, the majority of TnSeq studies published to date have used the lab-adapted reference strain, ATCC 19977 (25–27). One TnSeq study was conducted on a *massiliense* clinical isolate; however, the transposon library was not dense enough to quantify genetic requirement (28). Currently, there are no published studies investigating the functional diversity of *Mab* clinical isolates. To accomplish this, we conducted TnSeq on 21 *Mab* strains, generating triplicate libraries for 15 *abscessus* isolates and 6 single libraries of the *massiliense* isolates. Across the 21 strains, we identified 425 genes, representing ∼15% of the *Mab* core genome, as differentially required among isolates. We also identified genes whose differences in genetic requirements were sub-species, lineage, and isolate-specific. Finally, we identified pairs of genes whose genetic requirements were correlated, revealing previously undescribed genetic networks in *Mab*. Overall, we find that the genetic diversity in *Mab* clinical isolates is matched with significant functional genomic diversity as well.

## Results

### Characterization of *Mab* clinical isolates

Our lab obtained 25 *Mab* clinical isolates previously collected from patients in Taiwan and locally at Brigham and Women’s Hospital (BWH) (Supplemental Table 1). The clinical isolates represented a convenient sample of isolates to functionally characterize, given that they were isolated from multiple sites of infection, included both smooth and rough colony-morphology variants, and comprised both *abscessus* and *massiliense* subspecies. We passaged the clinical isolates only once to decrease selection of mutants under *in vitro* growth conditions. We then determined the genetic diversity of our clinical isolates through whole genome sequencing (WGS), followed by *de novo assembly* using ABySS (29) and annotation of the genomes using RAST (see Methods). The average number of nucleotides in the assemblies was 5103719 bp. Summary statistics of the WGS data are presented in Supplemental Table 2. A phylogeny of the 25 clinical isolates and five reference strains is shown in Figure 1A. The tree was generated using genome-wide SNP data while excluding SNPs from regions that showed evidence of a high degree of recombination, as determined by Gubbins (30). To put these isolates into a broader genomic context, they were incorporated into a global phylogeny constructed from 307 *Mab* isolates from Bryant *et al.* (11) using TNT (31). The resulting tree recapitulated the three subspecies of *Mab*, with our clinical isolates distributed across the entire tree (Figure 1B). About 60% of our isolates fell into clusters previously defined as dominant circulating clones (specifically *abscessus* DCC1 and *massiliense* DCC3) and are listed in Supplemental Table 1 (32).

**Figure 1:**
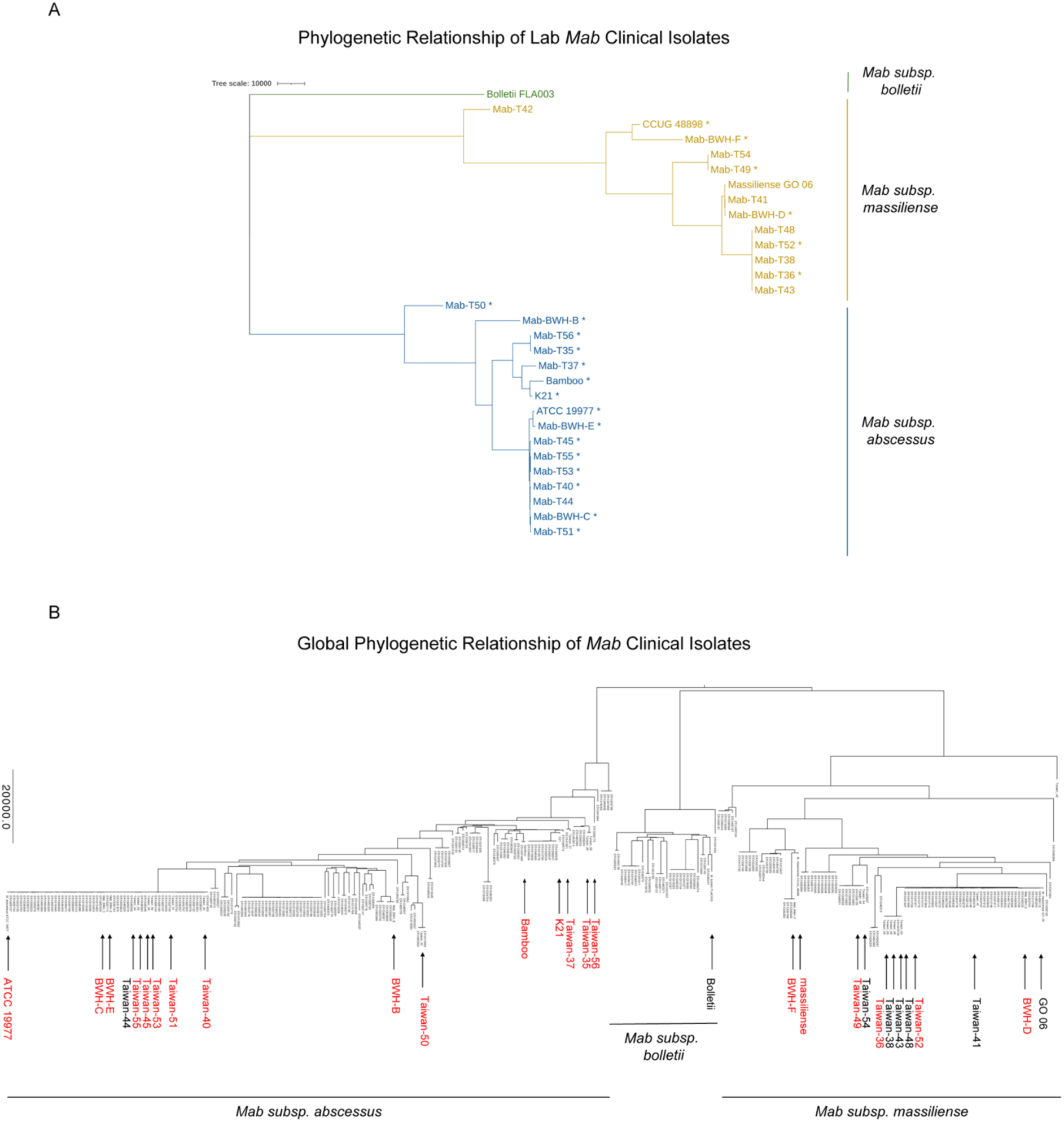
Phylogenetic Relationship of *Mab* Clinical Isolates. (A) Phylogenetic tree of 25 Mab clinical isolates and 5 reference strains. The tree is generated based on genome-wide SNPs, using Gubbins to filter out genomic regions potentially affected by recombination. Strains with transposon mutant libraries are marked with an ‘*’ **(B)** Global phylogenetic tree of *Mab* clinical isolates using genomes of 307 clinical isolates from Bryant *et al.*, 2016. Strains in the lab are labeled with an arrow. Red arrows indicate strains that have transposon-mutant libraries.

### Analysis of *Mab* core and accessory genome content

Next, we identified the clinical isolates’ core and accessory genome content using the Ptolemy pan-genome analysis software program (33). From our panel of *Mab* clinical isolates, we identified 15,352 gene clusters representing distinct open reading frames (ORFs) (Supplemental Table 3). Of these, ∼25% of the gene clusters were in the core genome, defined as present in at least 95% of the 30 isolates. The remaining ∼75% of gene clusters, or 11,507 ORFs, were in the accessory genome (Figure 2A). The annotated genomes contained ∼5200 genes, with accessory genes ranging from 997 to 1688. On average, the core genome comprised 75.6% of the total gene content in each strain, while the accessory genome accounted for 24.4%. These values are consistent with results published from previous studies (13,34). As expected, isolates more distantly related to the reference ATCC 19977 strain generally shared less accessory genome content (Figure 2B). The diversity of the accessory genome was remarkable, as evidenced by the number of isolates that contained distinct ORFs not present in any other isolates (Figure 2C). Of the 11,507 accessory genes identified, nearly 64% (n = 7361) were present in only one strain.

**Figure 2:**
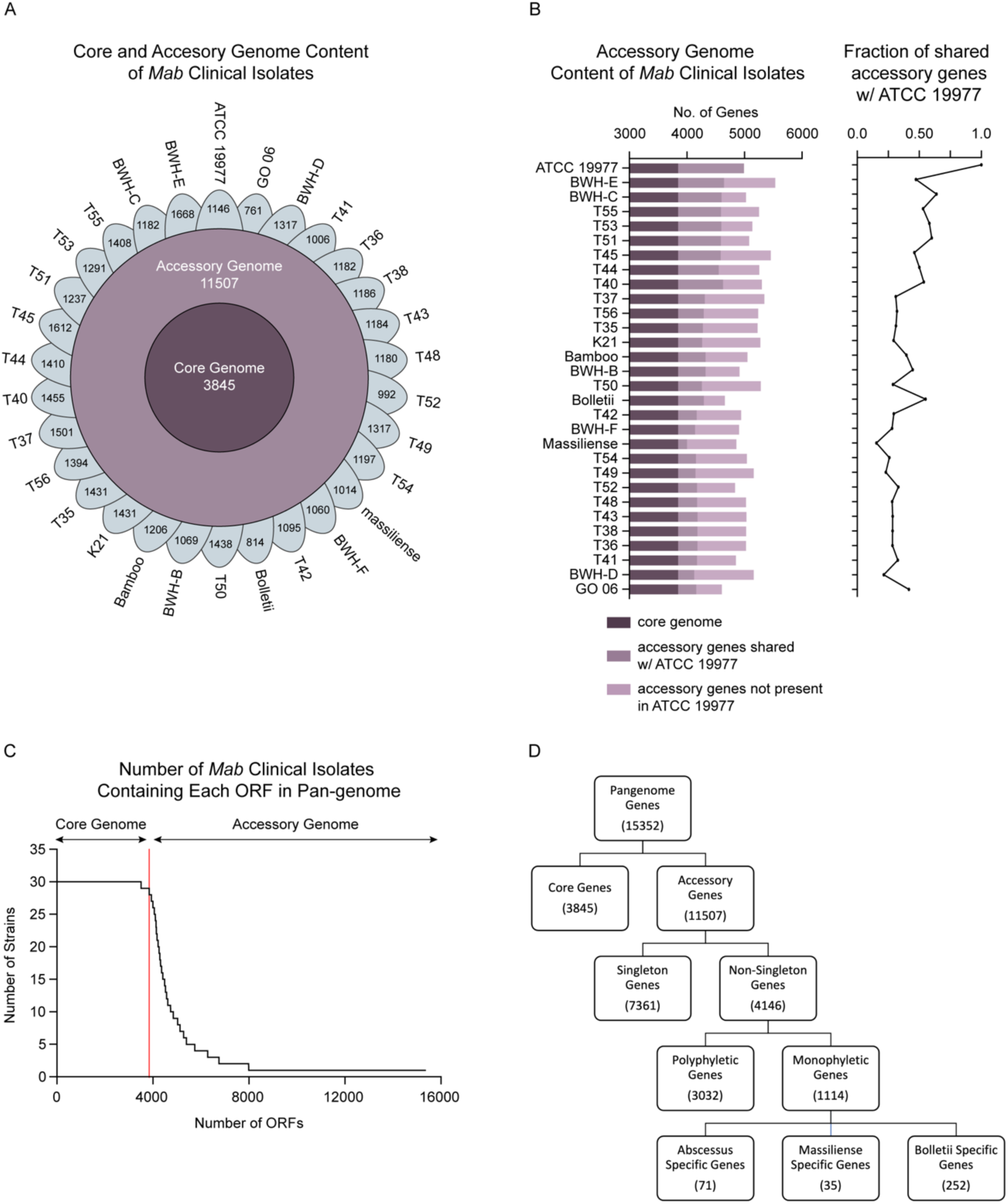
*Mab* has a large accessory genome. **(A)** Flower Pot diagram showing the breakdown of core and accessory genes in lab *Mab* clinical isolates. The outer leaflets represent the number of accessory genes in each clinical isolate. Strains are arranged counterclockwise based on the degree of similarity to the reference strain by core genome. **(B)** (*left*) Breakdown of accessory genome content in each clinical isolate. (*right*) Percentage of accessory gene content shared by the reference strain. **(C)** Number of ORFs present in the clinical isolates **(D)** Categorization of ORFs in the pan-genome.

The remaining 4147 genes were present in multiple (2–28) isolates. Of these, 1114 were monophyletic genes, meaning they were either inserted or deleted at one evolutionary time point and subsequently inherited by all the strains on that branch of the phylogenetic tree. The remaining 3210 accessory genes were polyphyletic. A subset of the monophyletic genes was sub-species specific and present in only *abscessus, massiliense,* or *boletti* isolates (Figure 2D).

### Transposon-sequencing of multiple *Mab* clinical isolates

Given the large genomic diversity present in *Mab*, we performed TnSeq across multiple clinical isolates to characterize the species’ functional diversity. The goal was to identify genes that were differentially required among the isolates to better understand the functional diversity in *Mab.* Specifically, we sought to identify a core set of essential genes in *Mab* clinical isolates and genes whose essentiality was sub-species or lineage-specific. To do this, we selected 21 isolates for TnSeq, which spanned the phylogenetic tree of *Mab*, including 15 *subsp. abscessus* and 6 *subsp. massiliense.* We then generated libraries in triplicate for the *abscessus* subsp. isolates, and single replicate libraries for the *massiliense* subsp. isolates. A schematic of the TnSeq experiment is presented in Supplemental Figure 1A. The transduction efficiency of the clinical isolates varied greatly, with the highest efficiencies observed in the two reference strains: ATCC 19977 and *massiliense* CCUG 48898 (hereafter referred to as *massiliense*) (Supplemental Figure 1B). After transduction, >150,000 independent mutants were harvested from each replicate, and sequencing libraries were prepared for next-generation sequencing. Of note, *subsp. bolletii* libraries were not generated in this study due to the absence of clinical isolates in the lab.

Sequencing revealed that nearly all the transposon libraries were well-saturated, with a median saturation of 59.5% (‘TA’ dinucleotide sites containing a transposon insertion). This range of saturation spanned 20% to 70.3% (Supplemental Figure 1C). Libraries with less than 33% saturation were excluded from further analysis. Previous TnSeq studies demonstrated that downstream analyses of gene essentiality performed well on libraries with saturation between 30%-80%. This analysis is also supported by the sensitivity analyses for HMM, resampling, and ANOVA presented in the Supplemental Materials. The summary statistics for each library are presented in Supplemental Table 4. To assess the reproducibility of our replicates, we compared the mean number of insertions in each gene between pairs of replicate libraries for each strain. We found that most replicates were well correlated with a median Pearson r^2^ value of 0.88 (Supplemental Table 5).

### HMM reveals essential genes in *Mab* clinical isolates

Using the Hidden Markov Model (HMM) algorithm (35), we determined the essentiality of genes in each isolate. The genes were categorized as essential, non-essential, growth defect, or growth advantage. Essential genes have no insertions in all or most of their TA sites. Genes classified as ‘growth defect’ are not essential, but their disruption causes growth impairment, resulting in suppressed transposon insertion counts. Conversely, when disrupted, genes classified as ‘growth advantage’ cause a fitness benefit relative to wild type, resulting in increased insertion counts. The essential genes of the reference strain, ATCC 19977, have been previously reported (25,27). We compared our predicted essential or growth-defect genes for ATCC 19977 to those in Rifat *et al.* (25) and found an overlap of 393 genes, showing high concordance between these two studies of *Mab* essentials (Supplemental Figure 2A). We also compared our predicted *Mab* essentials and growth-defect genes with those from the related mycobacterium *M. tuberculosis* (*Mtb* strain H37Rv) (36). There were 403 essential genes in common among 2278 genes with mutual orthologs, demonstrating substantial conservation between the two mycobacterial species (Supplemental Table 6). In total, there were 353 essential genes in the intersection of all 3 datasets (Supplemental Figure 2A). These genes were significantly enriched in key cellular pathways such as protein translation, amino acid metabolism, nucleotide metabolism, cell-cycle control, and cell wall synthesis (Supplemental Figure 2B). We then used the HMM to determine essential genes in all the clinical isolates. The essentiality call for each gene is listed in Supplemental Table 7. The average number of essential (ES) genes across the clinical isolates was 345, with a range of 281–391.

The combined number of essential and growth-defect (ES+GD) genes (avg ∼423) is consistent with essentiality analyses of other mycobacteria, where essential genes typically represent 10-15% of the genome (27). There were 259 pan-essential genes that were ES or GD across all 21 Mab isolates. These genes are listed in Supplemental Table 7. Chi-squared analysis was used to identify 17 clade-specific essential genes (Supplemental Figure 4), such as *mqo* (MAB_3159c, malate:quinone oxidoreductase), which is only essential in subsp. *massiliense*.

### ANOVA identifies additional differentially required genes

Next, we identified differentially required genes, defined as genes whose disruption causes varying degrees of growth defect in isolates. Traditionally, the ‘resampling’ method from TRANSIT is used to identify differentially required genes between two strains (35). We performed resampling between each clinical isolate and the reference ATCC 19977 strain, and the conditionally essential genes are listed in Supplemental Table 8. Resampling between pairs of isolates showed greater similarity (fewer conditionally essential genes) in comparisons within the same subspecies than those between subspecies (Supplemental Figure 5). These results gave us further confidence that our libraries were of sufficient quality for further analyses.

To compare the transposon insertion counts across all strains simultaneously, we used ANOVA, which identified 425 genes in the pan-genome (with adjusted P-value<0.05) that were differentially required across all 21 clinical isolates (Supplemental Table 9). For each gene, the log2-fold-change (LFC) of the insertion counts in each isolate was calculated relative to the grand mean across all isolates. These LFCs are depicted in a heatmap (Figure 3). Hierarchical clustering of the isolates revealed that the *massiliense* isolates (T36, T49, T52, BWH-D, BWH-F, massiliense) and the *abscessus* subspecies group together (Figure 3). Thus, the phylogenetic tree generated from TnSeq data roughly recapitulates the phylogeny generated from SNP differences (Figure 1A). The 425 differentially required genes represent 11% of the *Mab* core genome. Furthermore, 337 of these genes are non-essential in the reference ATCC 19977 genome (Figure 3B). These data reinforce the concept that genes can be differentially required based on genetic background (24).

**Figure 3:**
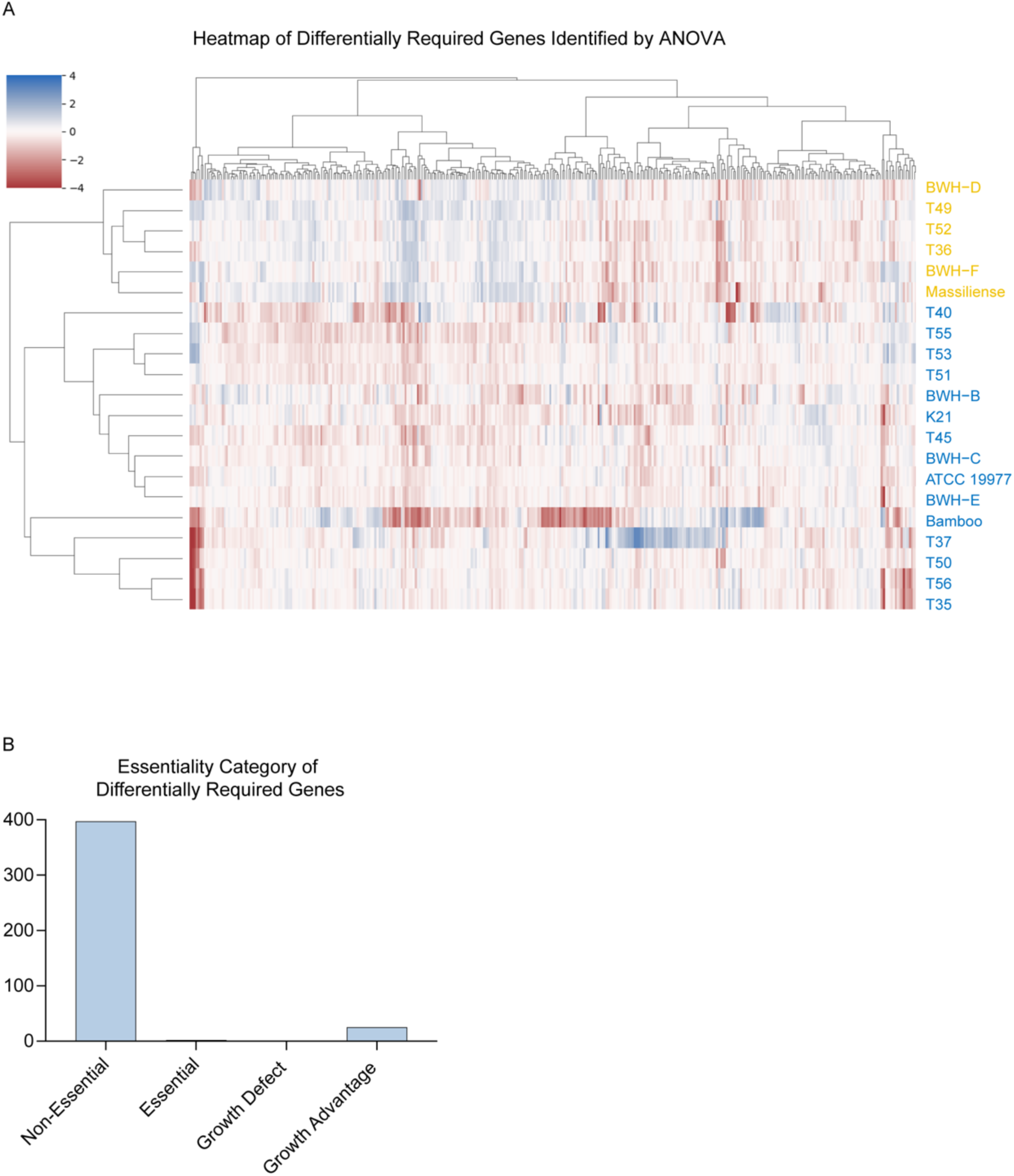
Summary of ANOVA results. **(A)** Heatmap depicting the log-2-fold-change (LFC) of insertion counts for the 425 differentially required genes in all isolates. **(B)** Essentiality category of the differentially essential genes in the ATCC 19977 reference strain.

### Gene essentiality varies by lineage and strain

To better understand the variance in the ANOVA data set, we used principal component analysis (PCA) to group the significant genes. The analysis showed principal components (PC) 1 & 2 explain ∼40% of the variation in the data (Figure 4A). When the isolates are clustered into three groups, they do so primarily by phylogenetic relationship, where cluster 1 is a sub-lineage of the *abscessus* subspecies, cluster 2 has the *abscessus* subspecies that are more closely related to the reference ATCC 19977 strain, and cluster 3 contains the *massiliense* subspecies. PCs 1 and 2 represent lineage-specific differences in gene requirement, while when PCs 3 and 4 are plotted, the phylogenetic clustering collapses (Figure 4B). The *Mab* phylogenetic tree is color-coded with the three corresponding clusters (Figure 4C). This analysis demonstrates that closely related isolates share genetic requirements and suggests a shared evolutionary origin of how these requirements arose.

**Figure 4:**
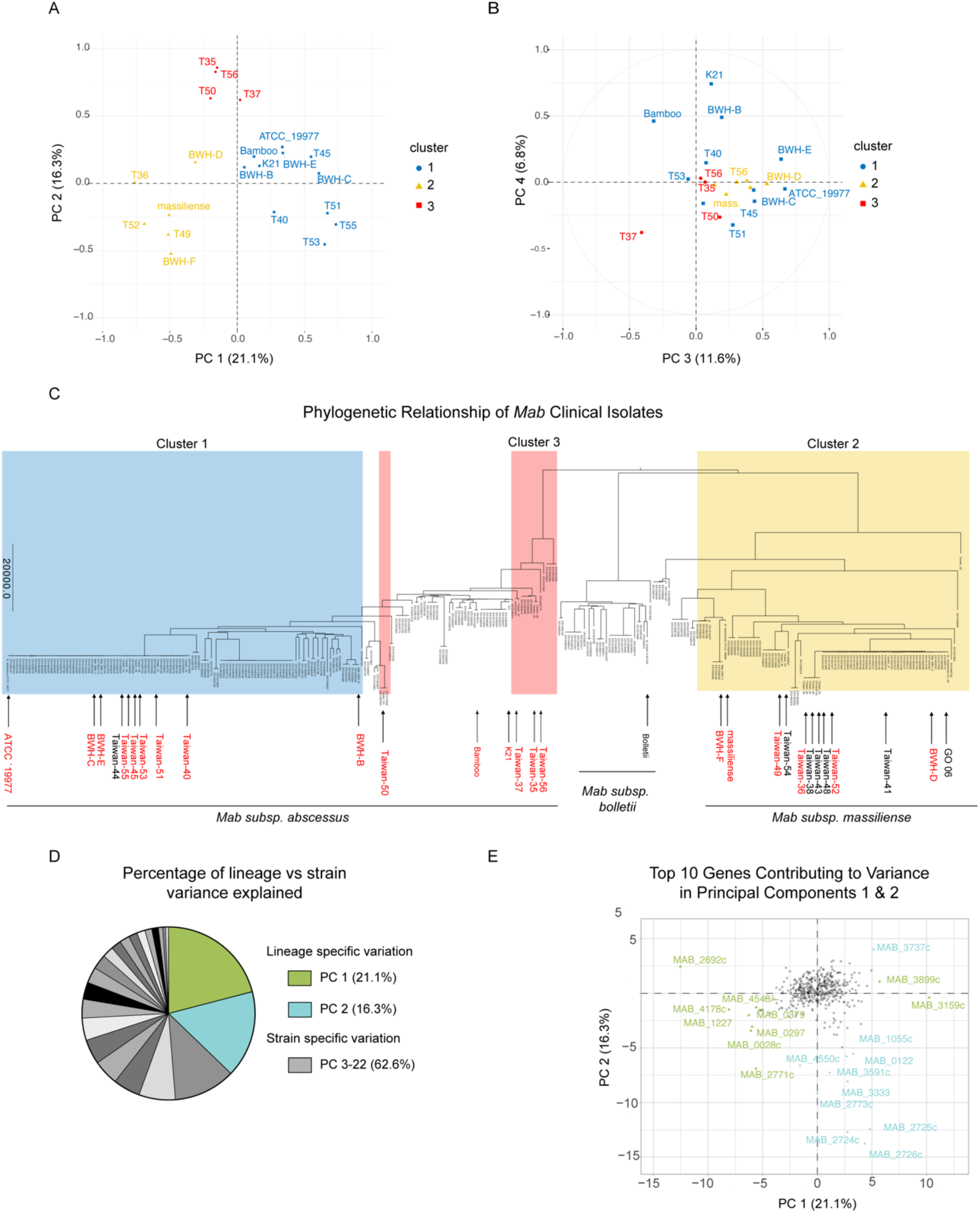
Principal Component Analysis of ANOVA Hits. **(A)** Principal components (PC) 1 & 2 of differentially required genes by ANOVA. **(B)** PC 3 & 4 **(C)** Phylogenetic tree of *Mab* clinical isolates shaded with rectangles highlighting clusters. **(D)** Percentage of variance explained by each of the principal components. **(E)** Plot depicting the top 10 contributing genes to the variance of PC 1 & 2.

The variation in PCs 1 and 2 only explains ∼40% of the variation in the dataset, corresponding to lineage-specific variation. In contrast, PCs 3-19 explain the remaining ∼60% of the variation and likely represent strain-specific variation or noise (Figure 4D). The top 10 genes contributing to the variance in PCs 1 and 2 are plotted in Figure 4E. The genes in PC 1 separate the two subspecies, while those in PC 2 separate Cluster 1 strains (T35, T37, T50, and T56) from the rest of the clinical isolates (Figure 4E).

### Identification of genes with lineage-specific essentiality

Next, we examined genes that exhibited lineage-specific differences in genetic requirement. We first identified genes that separated the reference strain from the clinical isolates. Two genes were the strongest differentiators: *MAB_4502c*, which encodes phosphoenolpyruvate carboxykinase (*pckA*), and *MAB_1327,* which encodes ferredoxin *(fdxC*) (Figure 5A). The reference strain requires *pckA* less than the clinical isolates, a difference that could be explained by laboratory adaptation of the reference strain.

**Figure 5:**
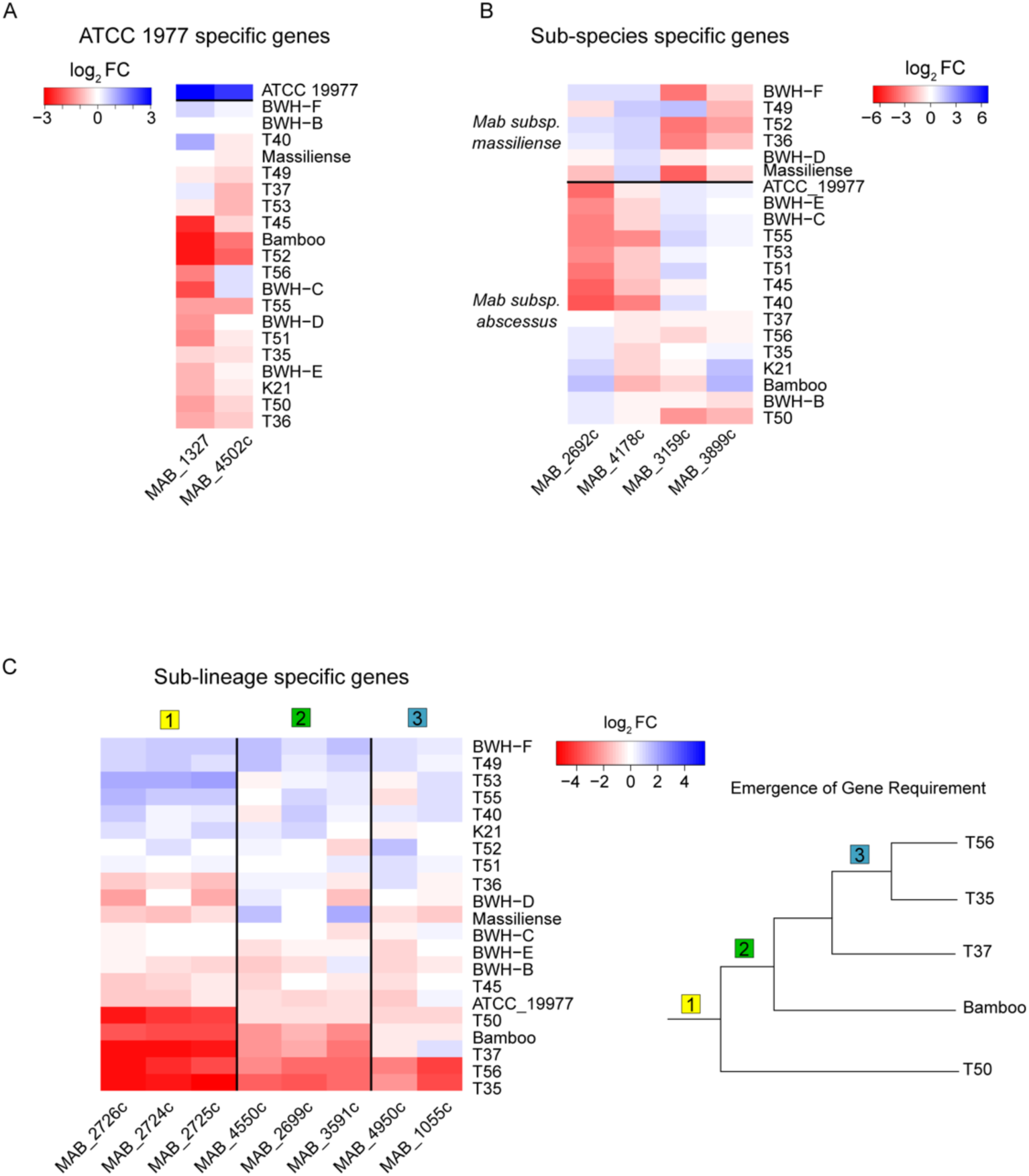
Sub-species and lineage-specific gene requirement. **(A)** Genes specific to the reference ATCC 19977 strain **(B)** Subspecies-specific genes that separate *abscessus* and *massiliense* subspecies. **(C)** (*left*) Genes that are differentially required in cluster 1. (*right*) Schematized phylogenetic relationship of strains in cluster 1.

Next, we analyzed the contributing genes in PC 1 that separate the two subspecies of *Mab abscessus* vs. *massiliense.* The top 4 genes are depicted in Figure 5B, while the top 20 genes that separate the subspecies are listed in Supplemental Table 10. We next inspected genes that separated strains in Cluster 1 (T35, T37, T50, and T56) from the rest of the clinical isolates (Figure 5C). The top 20 genes contributing to the variation in this cluster are listed in Supplemental Table 11.

The genes uniquely required in Cluster 1 can be split into three groups depending on how many strains share the level of gene requirement. For group 1, all four strains in the cluster share the same requirement for three genes in an operon, *MAB_2724c-MAB_2726c* (hypothetical genes of unknown function). Group 2 genes are shared only by T35, T37, and T56, which are more related to each other than T50. Finally, group 3 comprises genes shared by the two most closely related strains, T35 and T56. These data reveal genetic requirements emerging within a lineage. As strains diverge, their genomes change in genetic sequence and function, and genes that are otherwise dispensable in one strain can become more required in another. This is best illustrated by genes *MAB_4950c* and *MAB_1055c*, which are most strongly required in the two phylogenetically related isolates, T35 and T56 (Figure 5C).

### Correlated transposon insertion counts reveal functional genetic networks

To extract functional inferences from our TnSeq dataset, we looked for pairs of genes with correlated transposon insertion counts, which could reveal hidden functional genetic networks. To conduct this analysis, we selected the 751 unique ORFs with the most variability based on ANOVA (p-adj < 0.05) and other filters (see Methods). Of the 282,000 potential gene pairings, only 10,198 were significantly correlated, representing 3.6% of total potential pairs (Supplemental Table 12). In Figure 6A, we show three examples of gene pairs that are positively correlated. Pair 1 is genes *MAB_2725c x MAB_2726c*, whose insertion counts were correlated with an r^2^ of 0.92. The following two pairs, *MAB_2725c x MAB_3333* and *MAB_3246c x MAB_3333,* also have statistically significant correlated insertion counts. All these genes are correlated with each other and ostensibly form a network.

**Figure 6:**
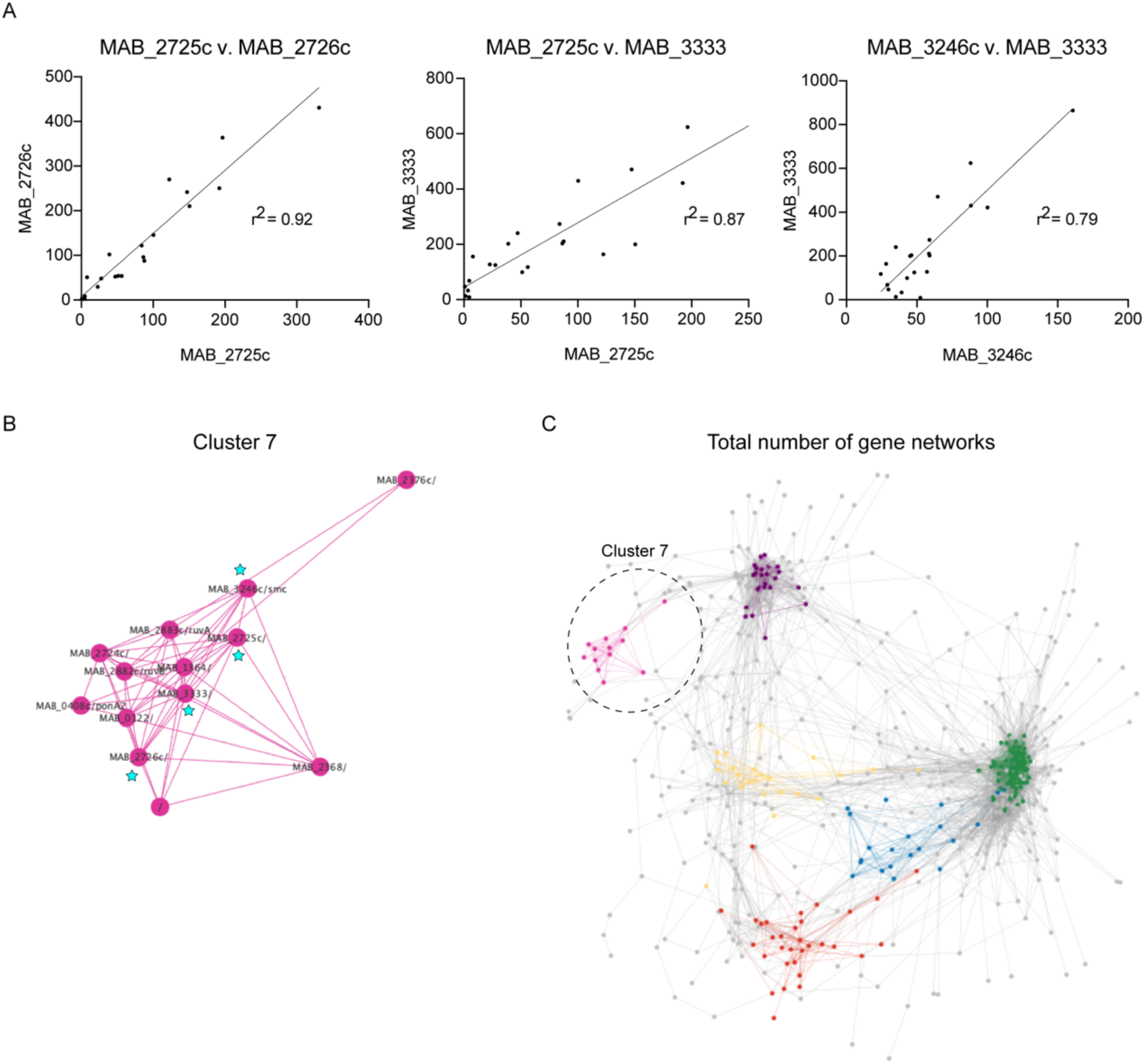
Network analysis of differentially required genes. **(A)** Correlation of transposon insertions between 3 gene pairs **(B)** Cluster 7 depicting 13 genes that are correlated with each other. Genes marked with blue stars are shown in panel A. **(C)** Network analysis depicting five additional clusters of 19 identified. Cluster 7 is highlighted by a dashed circle.

Based on these pairwise gene correlations, we performed network analysis with Cytoscape to turn these pairwise correlations into a network of interconnected genes. The network was organized into 19 clusters based on a greedy community identification algorithm (see Methods), so related genes with similar transposon insertion profiles were clustered. This analysis identified 19 clusters containing at least five genes (Supplemental Table 13). The genes profiled in Figure 6A are members of cluster 7 and are starred in the network (Figure 6B). Cluster 7 is comprised of 13 genes and includes the previously highlighted operon *MAB_2724c – MAB_2726* as well as 10 other genes, including *ruvA* and *ruvB*, which both encode Holliday junction ATP-dependent DNA helicases, and *MAB_3246c*, which encodes *smc*, segregation and condensation protein (Figure 6B). The complete network analysis with a few of the clusters highlighted is depicted in Figure 6C. Importantly, only comparative analysis of TnSeq data could reveal these genetic networks across multiple genetically diverse strains.

### Genes involved in central carbon metabolism are differentially required among clinical isolates

Resampling analysis unveiled several differentially required genes involved in carbon metabolism among the clinical isolates compared to the reference strain (Figure 7A). As noted above, *pckA* is more required in clinical isolates than ATCC 19977. Other examples include *mqo* (malate:quinone oxidoreductase), which is most required in T50 and BWH-F, and *pgi* (glucose-6-phosphate isomerase), which is most required in T35 and T56. To validate these differences, we employed CRISPRi/Cas9 to repress *pckA*, which is more required in nearly all the clinical isolates and is dispensable in ATCC 19977. We found that when *pckA* is repressed in ATCC 19977, there was no difference in CFUs between induction and non-induction plates as predicted by the TnSeq data (Figure 7B). However, when we repressed the enzyme in T35 and T45, we observed a 10X to 100X decrease in CFUs, indicating that *pckA* is more required in these clinical isolates. Notably, *pckA* was significantly more repressed in the reference strain than in the clinical isolates (Supplemental Figure 6A). The lack of reduction in CFUs in the reference strain despite stronger knockdown further supports *pckA’s* dispensability in ATCC 19977 relative to the tested clinical isolates.

**Figure 7:**
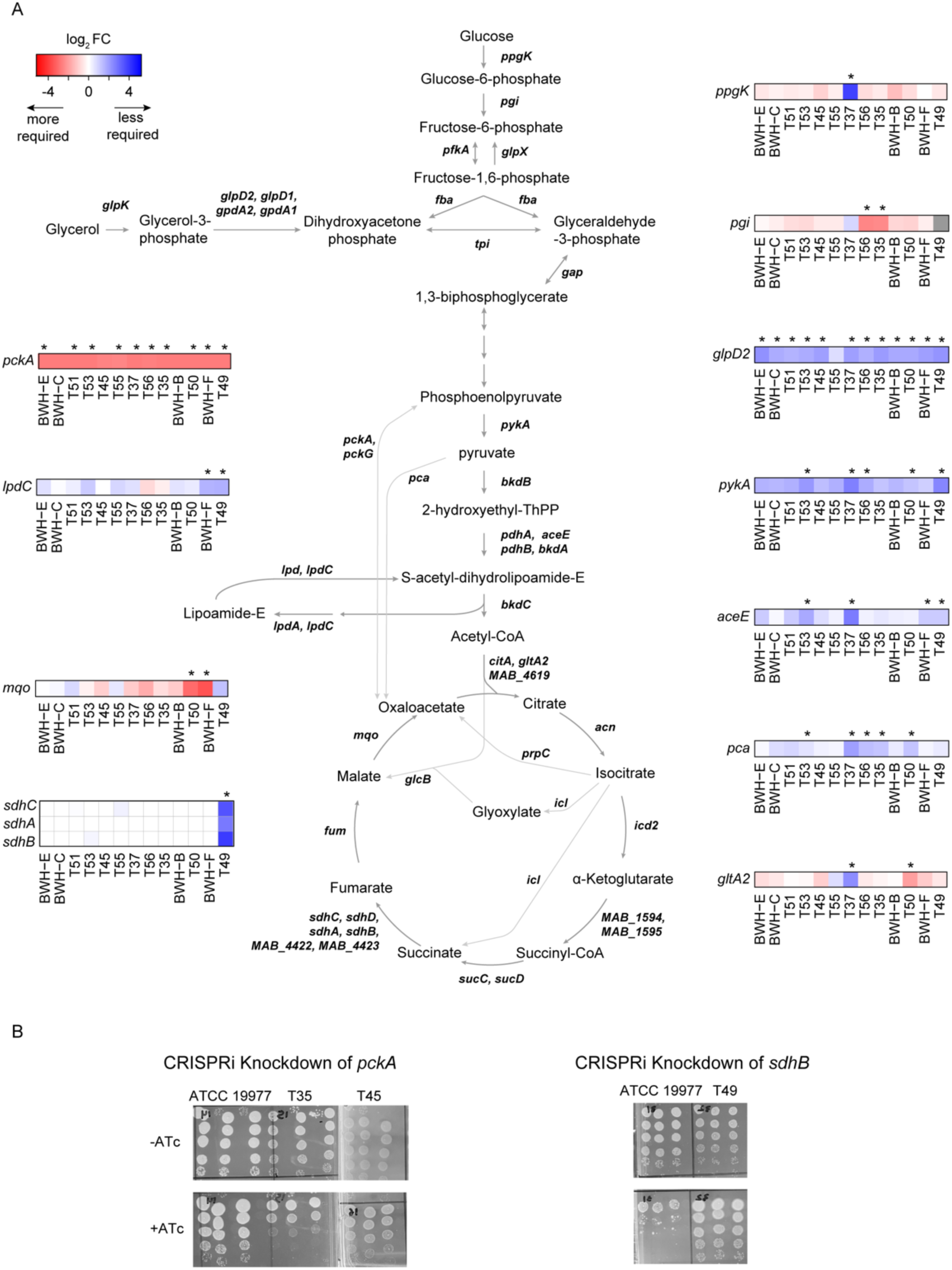
Genetic requirement differences in carbon metabolism genes among clinical isolates. **(A)** Schematic of genes in central carbon metabolism in *Mab*. All heatmaps showcase log-2-fold-change (LFC) relative to the reference strain. Genes with significant differences in LFC are highlighted with an “*” indicating a p-adj < 0.05. **(B)** CFU assay of *pckA* or *sdhB* clinical isolate knockdowns.

We also repressed *sdhB*, which was predicted to be required in ATCC 19977 and less required in the T49. Repressing *sdhB* in ATCC 19977 led to a 100X fold decrease in CFUs. However, when *sdhB* was repressed in T49, there was no difference in CFUs, confirming that *sdhB* is less required in T49 (Figure 7B). These differences in CFUs were observed with similar levels of *sdhB* knockdown (Supplemental Figure 6B). Altogether, these knockdown experiments validate our genetic requirement data and corroborate the functional differences in carbon metabolism between the clinical isolates.

## Discussion

While several groups have described the genomic diversity of *Mab* clinical isolates (13,16,17), to date, no studies have interrogated the functional consequence of this diversity. To investigate the functional diversity and genetic requirements of *Mab*, we performed TnSeq on 21 *Mab* clinical isolates spanning both the *abscessus* and *massiliense* sub-species. Among the 21 isolates, we identified 425 genes, or ∼11% of the *Mab* core genome as being differentially required among the isolates. This includes non-essential genes in the reference strain that became either fully essential or more required in some clinical isolates.

Applying TnSeq to a genetically diverse group of isolates enabled us to uncover functional differences among core and accessory genes. Using PCA and network analysis, we identified significant clusters of genes whose insertion patterns were correlated and whose functional requirements varied with phylogenetic position, either subspecies or lineage-specific. This data implied that *Mab* isolates have evolved to require a distinct subset of genes. Given that our genetic screen was conducted under *in vitro* conditions, passaging our transposon mutant libraries through more environmental or disease-relevant conditions would reveal additional variability in genetic requirement. Recently, Sullivan *et al.* passaged a transposon mutant library of clinical isolate T35 on an *in vitro* lung model and identified key genes responsible for establishing lung infection (37).

We also identified differences in gene requirement that were lineage-specific. Strains T35, T37, T50, and T56 represent one distinct lineage of the *abscessus* sub-species with genetic requirements unique to these strains. Notably, the operon *MAB_2724c-MAB_2726c* was uniquely required in this lineage and contributed 20% of the variance separating this sub-lineage from all the other clinical isolates. All three genes in the operon are annotated as hypothetical proteins (orthologous to Rv1479-81 in *Mtb*), and future studies are needed to understand why this operon is uniquely required in this subset of strains.

Interestingly, our study identified functional differences between the reference strain and the clinical isolates in genes related to carbon metabolism. Specifically, the gene *MAB_4502c* encodes the enzyme phosphoenolpyruvate carboxykinase, *pckA*, and is more required in nearly clinical isolates than the reference strain. *pckA* catalyzes the conversion of oxaloacetate to phosphoenolpyruvate, the first committed step of gluconeogenesis. The reduced requirement for *pckA* in ATCC 19977 is likely a consequence of the reference strain passaged on rich media and adapting to ample carbon sources for 70+ years (38).

This study raises an open question: what role does the accessory genome play in driving variability in genetic requirements among *Mab* strains? *Mab* is well-known to have a large accessory genome, which we confirmed using pan-genome analysis. Gene acquisition and gene loss through plasmid transfer, phage integration, duplication, or large-scale deletion likely influences genetic requirement and functional differences between strains. More work is needed to untangle the effects of these accessory genes’ presence and absence on the core genome’s essentiality. Nevertheless, this TnSeq dataset offers valuable insight into variable functionality among *Mab* strains and provides a foundation for future study of the biological and clinical consequences of genetic diversity of *Mab* clinical isolates.

## Methods

### Bacterial strains and culture conditions

*Mab* clinical strains were isolated from patients in Taiwan (T) and Brigham and Women’s Hospital (BWH), Boston, MA. Clinical isolates were streaked from patient stabs; single colonies were picked, cultured, and frozen for future use. Strains with transformed CRISPRi/Cas9 vectors to knockdown central carbon metabolism genes were built as described in (27). All strains were grown shaking at 37°C in liquid 7H9 media consisting of Middlebrook 7H9 salts with 0.2% glycerol, 0.85 g/L NaCl, and OADC (oleic acid, 5 g/L albumin, 2 g/L dextrose, 0.003 g/L catalase), and 0.05% Tween80. All *Mab* strains were plated on Middlebrook 7H10 agar supplemented with 10% OADC and 0.05% Tween80. Kanamycin 100 µg/ml was used for selection. All CRISPRi/Cas9 plasmids were induced with 500 ng/µl ATc.

### Whole Genome Sequencing of *Mab* Clinical Isolates

Genomic DNA from late-log phase cultures were extracted via a bead-beating and phenol-chloroform protocol described in Akusobi *et al.* (27), using an Illumina Rapid HiSeq 2500 instrument with 125+125 bp paired-end reads.

### Genome Assembly

Genome sequences for each isolate were assembled *de novo* from .fastq files using ABySS (v2.0) with default parameters (29). The assemblies had a mean total length of 5.14 Mbp (range: 4.86-5.54 Mbp) (Supplemental Table 2). To assemble core genomes, the contigs were mapped onto the *M. abscessus* ATCC 19977 (NC_010397.1) reference genome using Blastn and concatenated in syntenic order. The final genome length and depth of coverage are shown in Supplemental Table 2. To distinguish among *Mab* subspecies, the pattern of nucleotide substitutions was examined in *hsp65/groEL2* and rRNA (internal-spacer) loci (39).

### Genome Alignment and Annotation

Genome alignments were made using MUMMER v3.23 (40). To identify SNPs and indels, regions with genetic variation were aligned using the Needleman-Wunsch algorithm (41). For core genomes, the coordinates of open reading frames (ORFs) were adjusted for each isolate using a whole-genome alignment to the reference genome. Genes whose start or end coordinates were deleted in a clinical isolate were removed from the annotation. To produce unbiased annotations of each clinical isolate, we ran RAST (PATRIC CLI v1.039) on the contigs generated by ABySS (42). These annotations contain all predicted ORFs on all contigs.

### Phylogenetic Analysis

An alignment of all the core genomes was first generated to construct a phylogenetic tree incorporating all the *Mab* clinical isolates and reference strains. The included reference strains are listed in Supplemental Table 2, with sequencing data for *M. abscessus* str. K21 kindly provided by M. Gengenbacher. For all isolates, regions with insertions or deletions were filtered out, along with sites exhibiting low coverage (<10x), heterogeneity (<70% base-call purity), or repetition (where an overlapping window of 35 bp matches with at least 33/35 bp elsewhere in the genome). Finally, the remaining list of SNPs was extracted and used as input to RAxML (43) to generate the maximum likelihood phylogenetic tree, rooted using *M. bolletii* str. FLAC003 as an outgroup. Gubbins (30) was used to identify and filter out genomic regions (at least 500 SNPs) potentially affected by recombination, totaling <15% of the genome.

### Pan-Genome Analysis

The pan-genome analysis was performed on 30 Mab strains (25 clinical isolates and 5 reference strains) using Ptolemy v1.0 (33) to determine mappings of orthologs among ORFs based on homology and synteny. ABySS assemblies were input for each isolate and the ORFs from the RAST annotation. The output was used to generate a spreadsheet with all “pan-ORFs” (15,352 ortholog clusters) as rows and individual ORF IDs in columns. This data is provided in Supplemental Table 3, which also contains a count of how many of the 30 strains had a copy of each pan-ORF, which was used to differentiate core genes (genes with orthologs in >95% of isolates) from accessory genes (present in 1-28 isolates).

### Generation of Transposon Libraries

High-density transposon libraries of selected *Mab* clinical isolates were generated using the ϕMycoMarT7 phage described in Akusobi *et al.* (25–27). Libraries were generated in biological triplicate for 15 clinical isolates, while single replicate libraries were generated for 6 isolates (see Supplemental Table 4). K21 was a generous gift from Thomas Dick, Meridian Health, Hackensack, NJ.

Libraries were selected for 4 days on 7H10 agar supplemented with 10% OADC, 0.5% glycerol, 0.05% Tween80, and 100 µg/ml kanamycin. Libraries of >150,000 mutants were harvested and stored in 7H9 + 10% glycerol at −80°C for future use. Transposon-sequencing libraries were prepared following the protocol described in (44). The TnSeq samples were sequenced on an Illumina Rapid HiSeq 2500 instrument with 125+125 bp paired-end reads.

### TnSeq Analysis

The .fastq files obtained from sequencing were pre-processed using TPP in Transit (35). The reads were then mapped to the syntenic versions of each genome (aligned to ATCC 19977) and the raw Abyss assemblies (all contigs) of each isolate. The median library saturation was 58.4% (19.0-72.6%) (Supplemental Table 4). BWH-B-2, BWH-C-1, and all T44 strains were excluded from further analyses due to low saturation or total insertion count.

The Hidden Markov Model (HMM) in Transit was run to identify essential and non-essential genes in each isolate (Supplemental Table 7). Furthermore, the genes were evaluated by a confidence score based on the consistency of the insertion statistics with the posterior distribution of the called essentiality state (see Supplemental Material). Low-confidence genes (<0.20) were filtered out of the analyses. The ‘resampling’ method in Transit was used to identify conditionally essential genes in each isolate relative to ATCC 19977. p-values were adjusted for multiple tests using the Benjamini-Hochberg procedure (45) for an FDR of 5%. Significant genes are those with P-adj < 0.05 (Supplemental Table 8). ANOVA analysis was then applied to the pan-ORFs to identify genes with significant variability of insertion counts across strains. The normalized counts at each TA site for each gene were tabulated and averaged at the strain level, excluding genes with mean < 20. Then one-way ANOVA was used to calculate the F-statistic and P-value, adjusted for multiple testing as above.

### Genetic Interactions and Network Analysis

A correlation analysis of mean transposon insertion counts was performed for gene pairs across all *Mab* strains. Genes with significant variability by ANOVA (P-adj < 0.05) and insertion count < 20 or Z-max > 8 (used to filter out outliers) were selected, resulting in 751 varying genes. A Pearson correlation coefficient was calculated for gene pairs that were jointly present in at least 10 strains. This analysis identified 10,198 pairs of genes with a correlation of > 0.7 and adjusted p-value < 0.05. A network diagram based on these correlations was generated in Cytoscape v3.9.1 using the Edge-weighted Spring-embedded Layout.

To extract clusters of genes with similar TnSeq profiles, a network community analysis was performed based on a greedy algorithm. The degree of each gene was calculated, reflecting the number of genes correlated to it with cc > 0.7. The gene with the highest degree represented the most “central” gene in the network and was required to be present in at least 15 strains. We extracted the cluster of genes connected to the most central gene as the first cluster. To extract subsequent clusters, we identified the next most central gene whose neighbors did not overlap more than 20% with genes in previously selected clusters. The process was iterated until clusters contained fewer than 5 genes.

### RNA Extraction and RT-PCR

For each strain, cultures were grown in biological triplicate to mid-log phase, diluted back in +/-500 ng/ml ATc, and grown for 18 hr to achieve target knockdown. Afterward, 2 OD_600_ equivalents of cells from each culture were harvested by centrifugation, resuspended in TRIzol (Thermo Fisher), and lysed by bead beating (Lysing Matrix B, MP Biomedicals). RNA extraction and subsequent RT-PCR experiments were performed as described by Akusobi et al. (27).

---

**Supplemental Figure 1:**
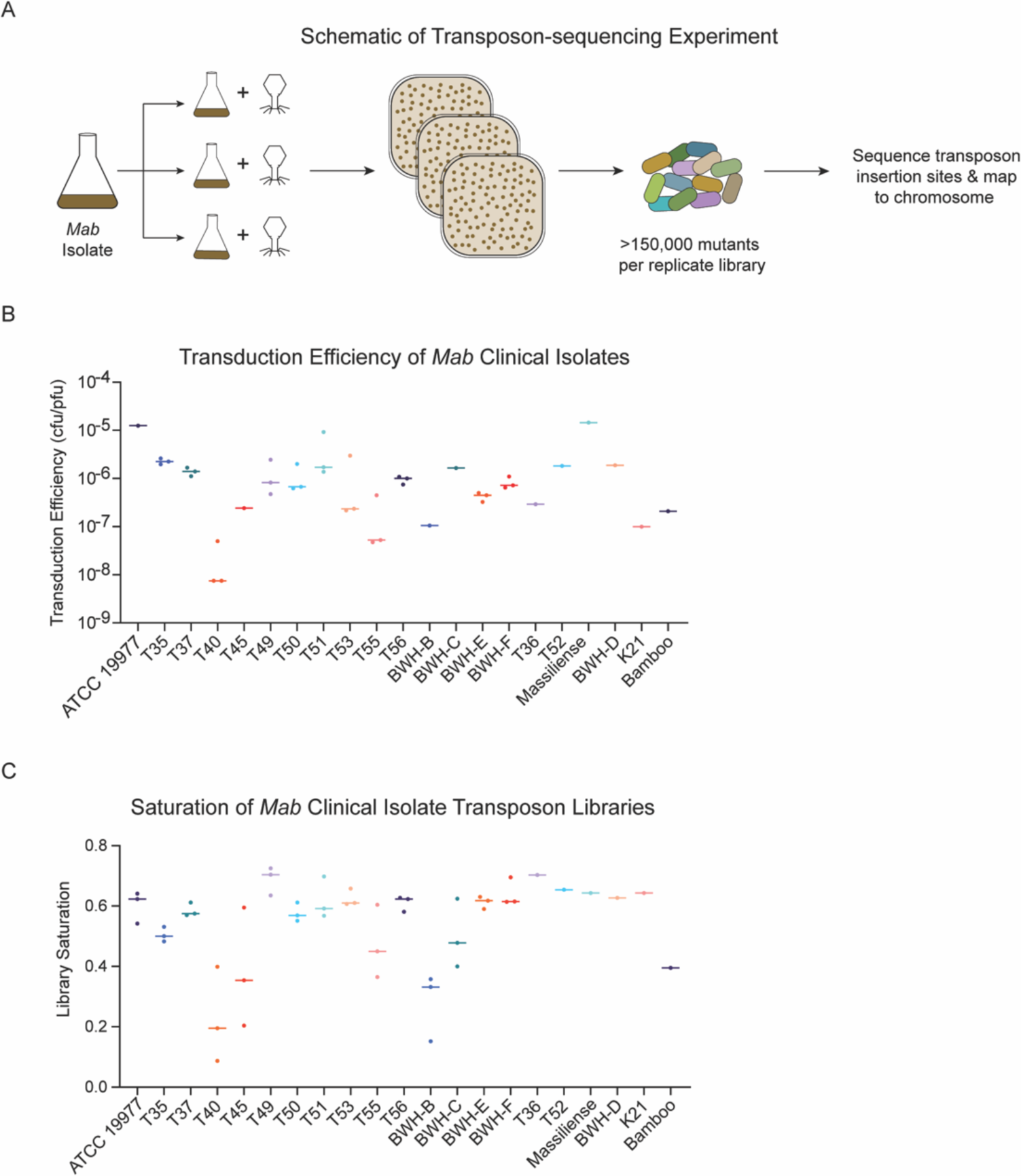
Summary of TnSeq data. **(A)** Schematic of TnSeq experiment. **(B)** Transduction efficiency (CFU/PFU) of *Mab* clinical isolates. (**C)** Percent saturation of TnSeq libraries. The dotted line demarcates 30% library saturation used as a threshold for study inclusion.

**Supplemental Figure 2:**
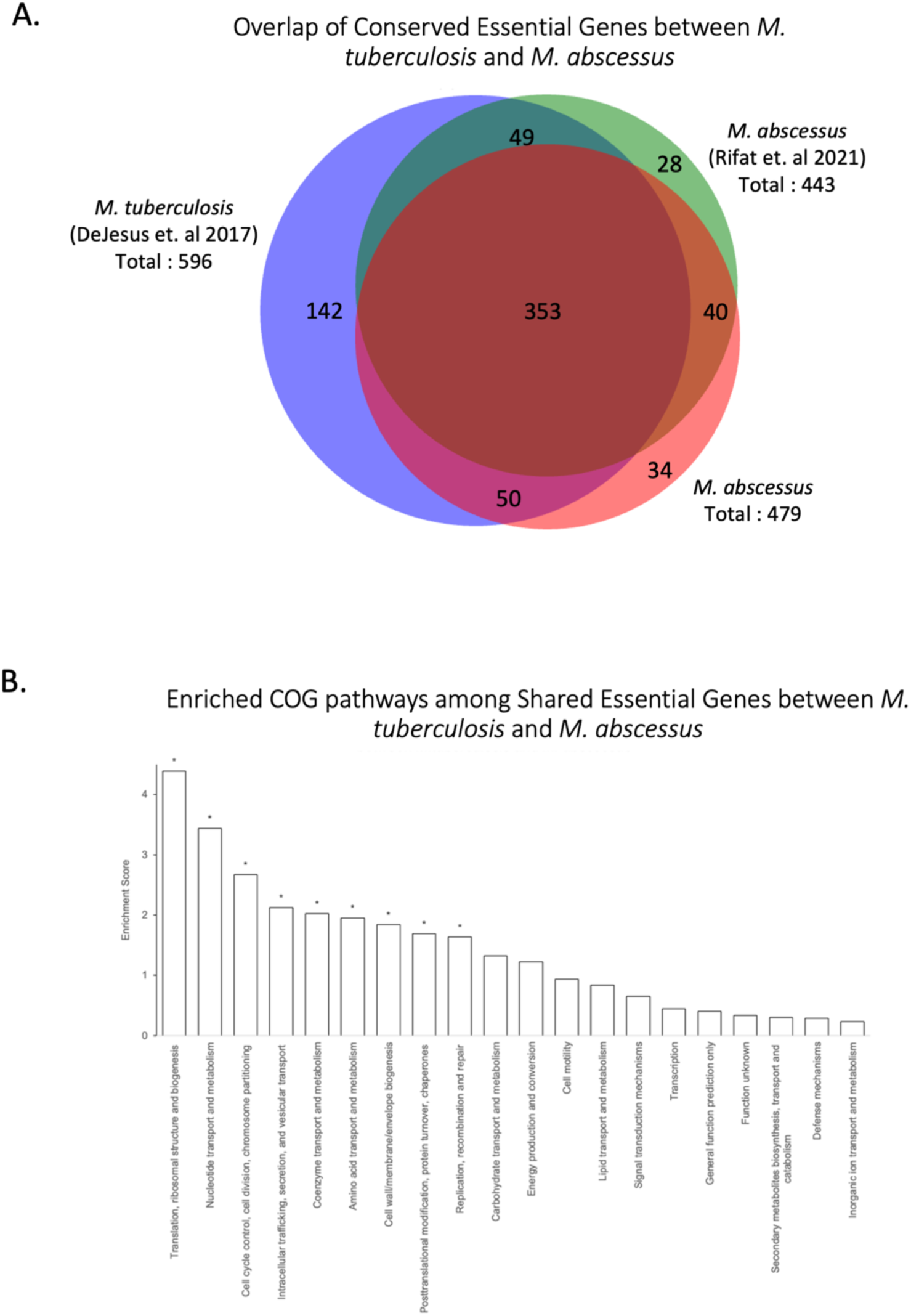
Comparison of *M. tuberculosis* and *M. abscessus* essential genes. **(A)** Overlap of conserved essential genes between *Mtb* and *Mab.* **(B)** Enriched COG pathways of shared essential genes between *Mtb* and *Mab*. Pathways demarcated with an “*” indicate p-adj < 0.05.

**Supplemental Figure 3:**
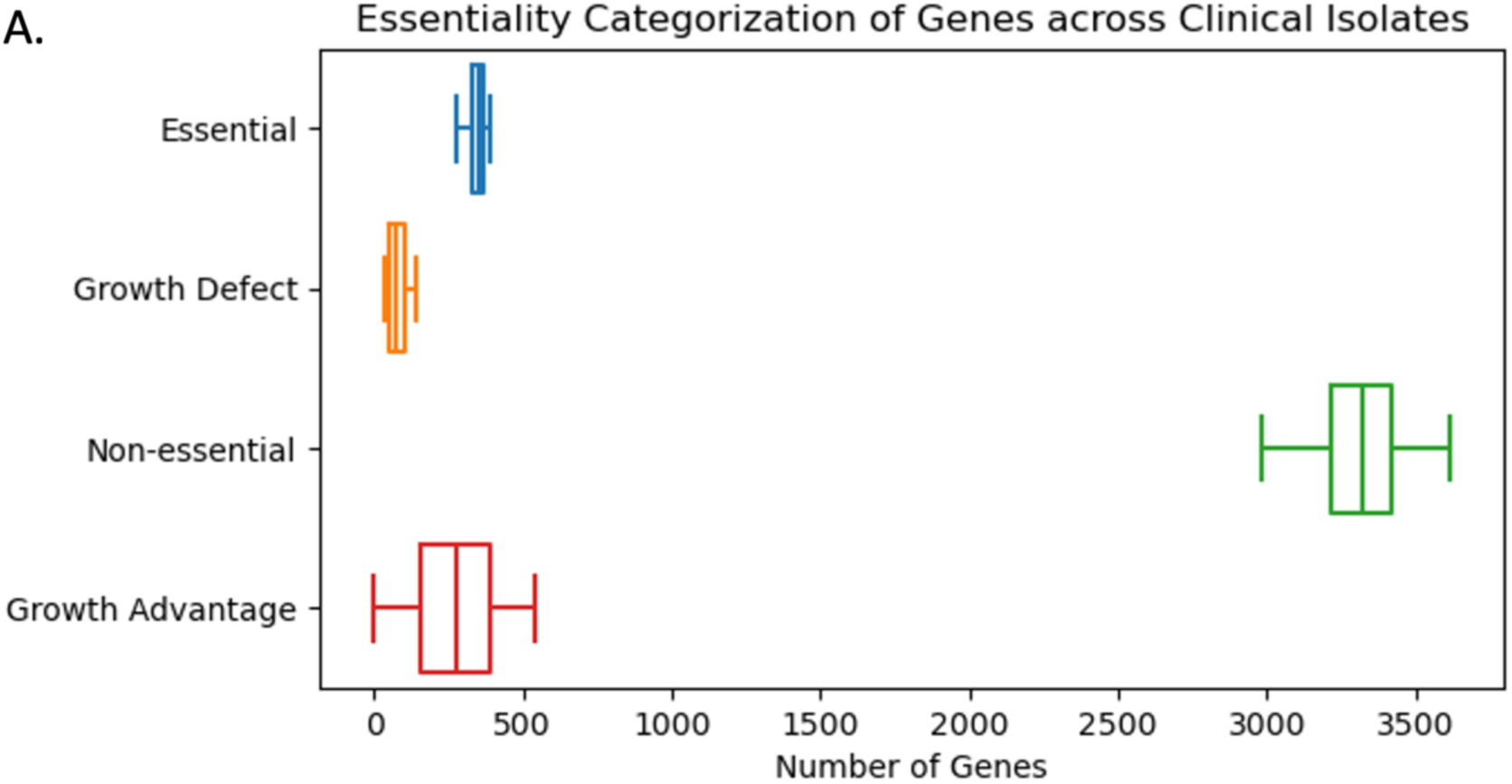
Summary of gene essentiality category across *Mab* clinical isolates.

**Supplemental Figure 4:**
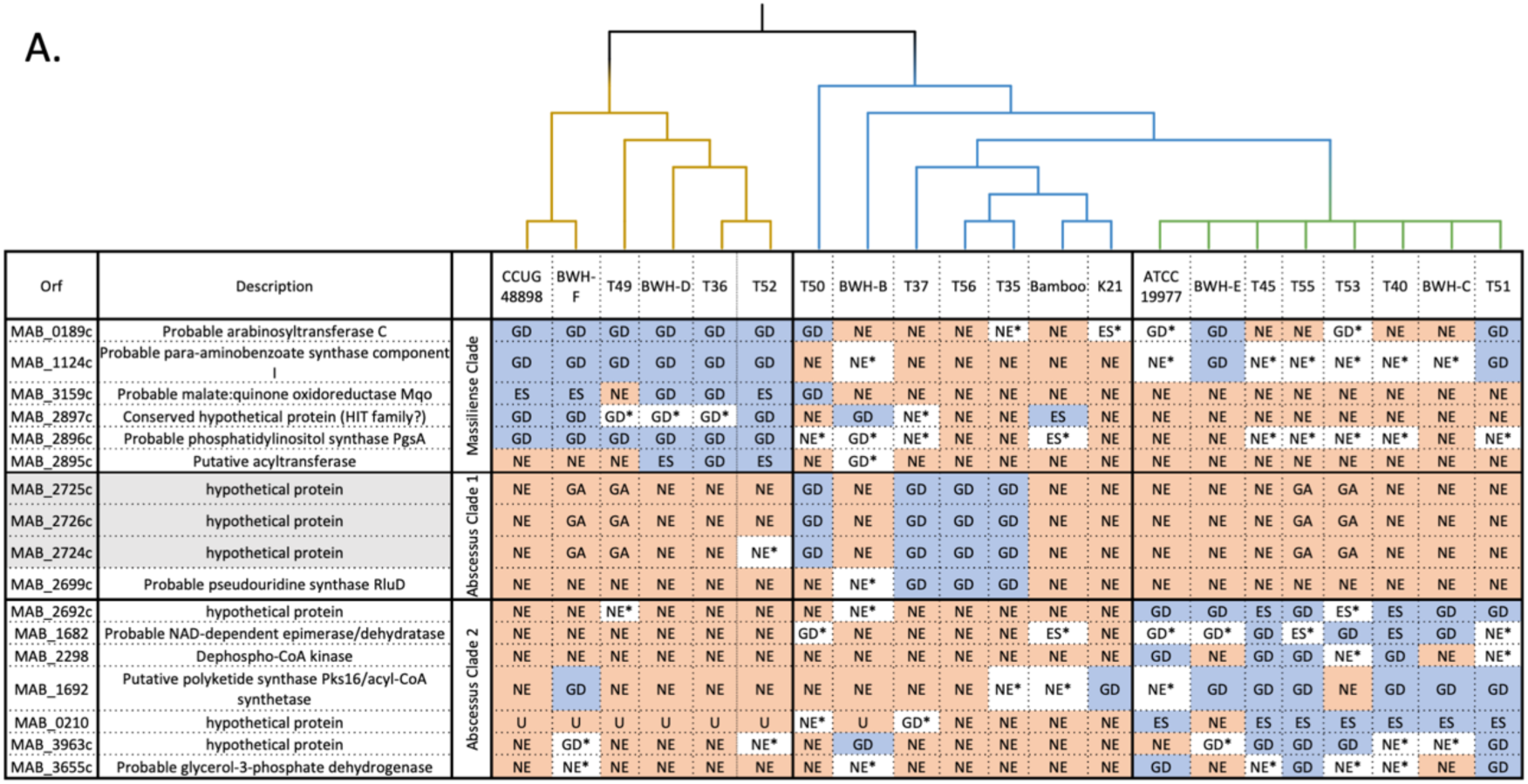
Analysis of shared essential genes in *Mab* clinical isolates. Analysis of clade-specific essentials in *Mab* clinical isolates. The HMM calls of the clade-specific genes across all 21 *Mab* isolates (ignoring genes with low-confidence calls, marked with an asterisk). The phylogeny of the 21 isolates is depicted above with the *massiliense* clade (includes all *massiliense* isolates) in yellow, the *abscessus* clade 1 in blue, and *abscessus* clade 2 in green. The highlighted gray genes are seen in Figure 5, depicting lineage-specific gene essentiality.

**Supplemental Figure 5:**
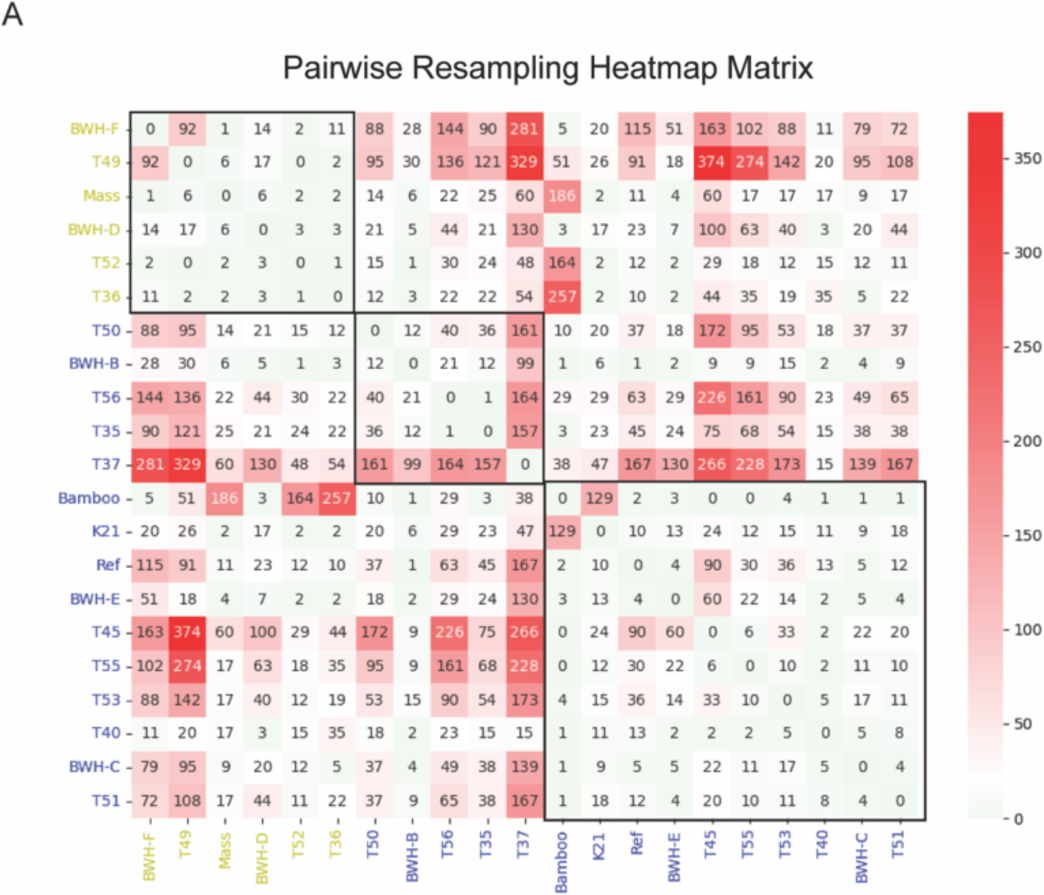
Pairwise Resampling Heatmap Matrix. Each number in comparison represents the differentially required genes between two clinical isolates. Top left corner represents subsp. *massiliense – massiliense* comparisons. Bottom right comparisons represent subsp. *abscessus – abscessus* comparisons with strains more closely related to the ATCC 19977 reference strain.

**Supplemental Figure 6:**
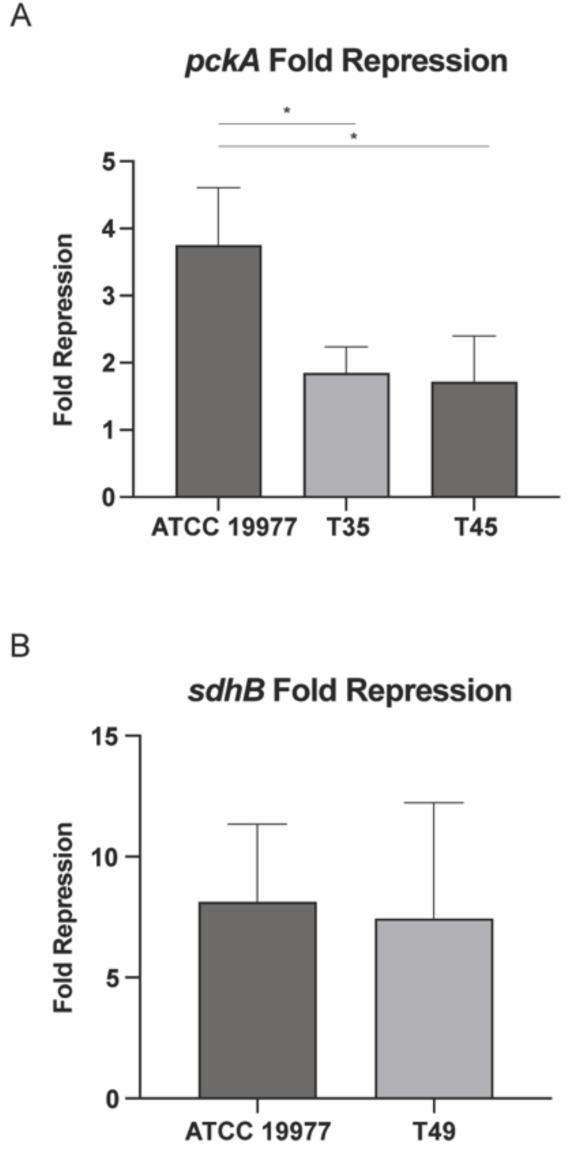
Fold Repression of *pckA* and *sdhB.* (A). Fold repression of *pckA* as measured by RT-qPCR. “*” signifies p < 0.05 by student’s t-test. **(B)** Fold repression of *sdhB* as measured by RT-qPCR.

