## SupplementalMaterial for "Transposon-sequencing across multiple *Mycobacterium abscessus* isolates reveals significant functional genomic diversity among strains"

### Supplemental Material

#### Functional Analysis of Conserved Essential Genes between *Mtb* and *Mab*

As illustrated in the Venn diagram in Figure SM1, there is a significant overlap in essential and growth defect HMM calls in Akusobi et al (for the ATCC 19977 reference strain) and those reported in Rifat et al [1]. Among the 2278 Mab genes with mutual orthologs in *M. tuberculosis* [2, 3], 353 genes are essential in *M. abscessus* (both datasets) and *M. tuberculosis* (see Supplemental Table 6).

Pathway analysis (Fisher's Exact Test, FDR-adjusted p-value<0.05) using COG pathways [4] of these 353 conserved essentials found significant enrichment of 9 pathways including translation, cell-wall biogenesis, nucleotide metabolism, etc. These data suggest that conserved essential genes are involved in key housekeeping functions. These include expected house-keeping genes such as subunits of the ribosome and RNA polymerase.

Within the 21 Mab isolates (spanning *subsp. abscessus* and *massiliense*), 259 pan-essential genes were identified ES or GD across all 21 isolates. The calls of each of the isolates along, with a summary can be seen in Supplemental Table 7.

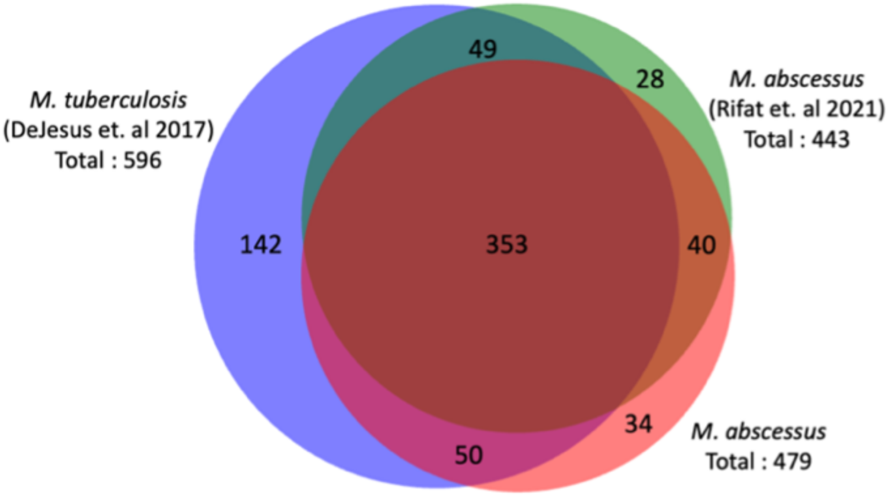


**Figure SM1. Overlap of conserved essentials in *M. tuberculosis H37Rv* and *M. abscessus* ATCC 19977.** There is a significant overlap between the shared essentials in Akusobi et al (for the ATCC 19977 reference strain) and those reported in Rifat et al [1], illustrated in the Venn diagram above. The orthologs of most of these *Mab* essential genes are also essential in *M. tuberculosis* [2, 3]. Among the 2278 genes with mutual orthologs (see Supplemental Table 6), 353 genes are essential in *M. abscessus* (both Rifat et al and Akusobi et al datasets) and *M. tuberculosis*.

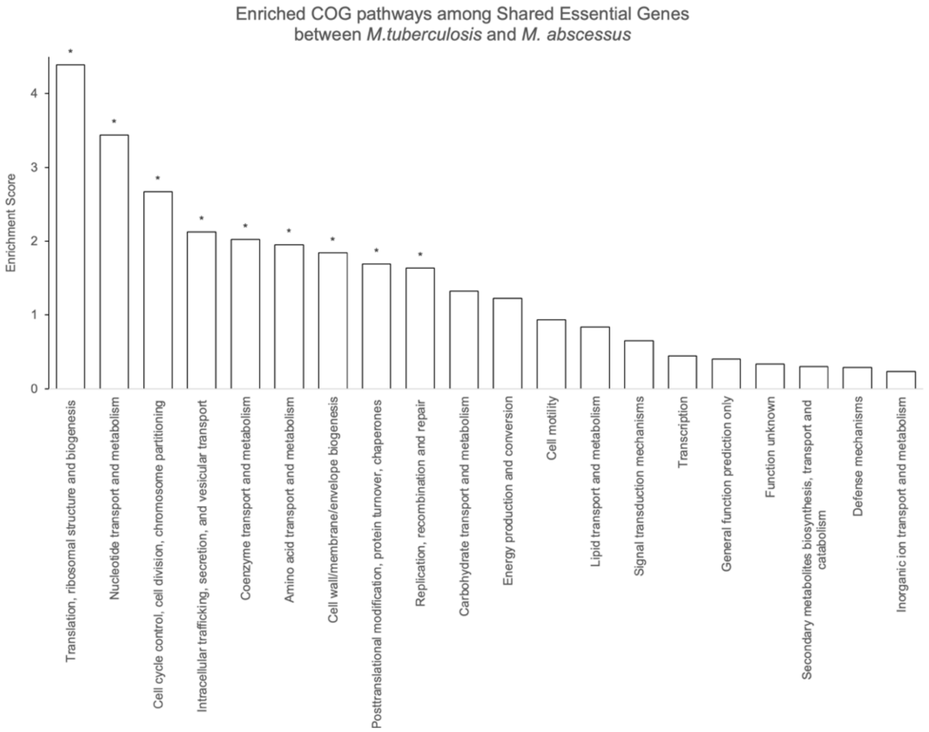


**Figure SM2. Enriched COG pathways among essential genes shared between *Mtb* and *Mab*.** Pathway analysis (Fisher's Exact Test, FDR-adjusted p-value<0.05) using COG pathways [4] of the 353 conserved essential genes among all 3 datasets (2 species) found significant enrichment of 9 pathways, including DNA replication, protein translation, cell-wall biogenesis, nucleotide metabolism, etc. The enrichment of these pathways suggests, unsurprisingly, that the conserved essential genes are involved in key cellular housekeeping functions.

### Functional Analysis of Conserved Essential Genes among *Mab* Clinical Isolates

#### Identifying Pan-essential and Clade-Specific Genes

Based on the phylogenetic tree (Figure 1A), the isolates divide into three clades:

1. *massiliense* clade (aligns with DCC3): CCUG 44898, BWH-F, T49, BWH-D, T36, T52
2. *abscessus* clade 1: T50, BWH-B, T56, T35, T37, Bamboo, K21
3. *abscessus* clade 2 (aligns with DCC1): ATCC_19977, BWH-E, T40, BWH-C, T51, T45, T55, T53

For each gene and each clade, we calculated a 2 x 2 contingency matrix between the variables representing clade and essentiality. A chi-squared test was then performed to evaluate the p-value of the gene's clade-specific association. We found 6 genes specific to the *massiliense* clade, 4 genes specific to *abscessus* clade 1 and 7 genes specific to *abscessus* clade 2. The clade-specific genes are shown in Figure SM3. An example of a clade-specific essential gene *mqo* (malate:quinone oxidoreductase, MAB_3159c), a gene that is uniquely essential in the *massiliense* clade (blue cells) but not the two clades. Also, disruption of *pks16* (MAB_1692) appears to cause a growth defect in *abscessus* clade 2 (blue cells), but not in the other two clades.

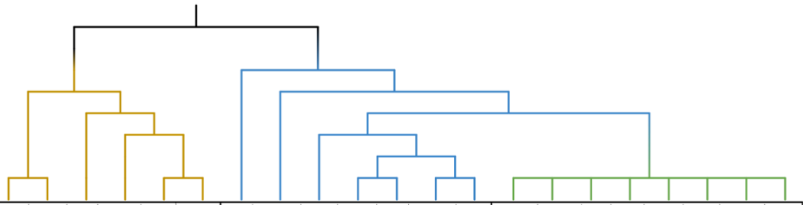

| Orf | Description | | CCUG 48898 | BWH-F | T49 | BWH-D | T36 | T52 | T50 | BWH-B | T37 | T56 | T35 | Bamboo | K21 | ATCC 19977 | BWH-E | T45 | T55 | T53 | T40 | BWH-C | T51 |
|---|---|---|---|---|---|---|---|---|---|---|---|---|---|---|---|---|---|---|---|---|---|---|---|
| MAB_0189c | Probable arabinosyltransferase C | Massiliense Clade | GD | GD | GD | GD | GD | GD | GD | NE | NE | NE | NE* | NE | ES* | GD* | GD | NE | NE | GD* | NE | NE | GD |
| MAB_1124c | Probable para-aminobenzoate synthase component I | | GD | GD | GD | GD | GD | GD | NE | NE* | NE | NE | NE | NE | NE | NE* | GD | NE* | NE* | NE* | NE* | NE* | GD |
| MAB_3159c | Probable malate:quinone oxidoreductase Mqo | | ES | ES | NE | GD | GD | ES | GD | NE | NE | NE | NE | NE | NE | NE | NE | NE | NE | NE | NE | NE | NE |
| MAB_2897c | Conserved hypothetical protein (HIT family?) | | GD | GD | GD* | GD* | GD* | GD | NE | GD | NE* | NE | NE | ES | NE | NE | NE | NE | NE | NE | NE | NE | NE |
| MAB_2896c | Probable phosphatidylinositol synthase PgsA | | GD | GD | GD | GD | GD | GD | NE* | GD* | NE* | NE | NE | ES* | NE | NE | NE | NE* | NE* | NE* | NE* | NE | NE* |
| MAB_2895c | Putative acyltransferase | | NE | NE | NE | ES | GD | ES | NE | GD* | NE | NE | NE | NE | NE | NE | NE | NE | NE | NE | NE | NE | NE |
| MAB_2725c | hypothetical protein | Abscessus Clade 1 | NE | GA | GA | NE | NE | NE | GD | NE | GD | GD | GD | NE | NE | NE | NE | NE | GA | GA | NE | NE | NE |
| MAB_2726c | hypothetical protein | | NE | GA | GA | NE | NE | NE | GD | NE | GD | GD | GD | NE | NE | NE | NE | NE | GA | GA | NE | NE | NE |
| MAB_2724c | hypothetical protein | | NE | GA | GA | NE | NE | NE* | GD | NE | GD | GD | GD | NE | NE | NE | NE | NE | GA | GA | NE | NE | NE |
| MAB_2699c | Probable pseudouridine synthase RluD | | NE | NE | NE | NE | NE | NE | NE | NE* | GD | GD | GD | NE | NE | NE | NE | NE | NE | NE | NE | NE | NE |
| MAB_2692c | hypothetical protein | Abscessus Clade 2 | NE | NE | NE* | NE | NE | NE | NE | NE* | NE | NE | NE | NE | NE | GD | GD | ES | GD | ES* | ES | GD | GD |
| MAB_1682 | Probable NAD-dependent epimerase/dehydratase | | NE | NE | NE | NE | NE | NE | GD* | NE | NE | NE | NE | ES* | NE | GD* | GD* | GD | ES* | GD | ES | GD | NE* |
| MAB_2298 | Dephospho-CoA kinase | | NE | NE | NE | NE | NE | NE | NE | NE | NE | NE | NE | NE | NE | GD | NE | GD | GD | NE* | GD | NE | NE* |
| MAB_1692 | Putative polyketide synthase Pks16/acyl-CoA synthetase | | NE | GD | NE | NE | NE | NE | NE | NE | NE | NE | NE* | NE* | GD | NE* | GD | GD | GD | NE | GD | GD | GD |
| MAB_0210 | hypothetical protein | | U | U | U | U | U | U | NE* | U | GD* | NE | NE | NE | NE | ES | NE | ES | ES | ES | ES | ES | ES |
| MAB_3963c | hypothetical protein | | NE | GD* | NE | NE | NE | NE* | NE | GD | NE | NE | NE | NE | NE | NE | GD* | GD | GD | GD | NE* | NE* | GD |
| MAB_3655c | Probable glycerol-3-phosphate dehydrogenase | | NE | NE* | NE | NE | NE | NE | NE | NE* | NE | NE | NE | NE | NE | GD | NE | NE* | GD | NE* | NE* | NE | GD |

**Figure SM3**. **Analysis of clade specific essentials in *Mab* clinical isolates**. The HMM calls of the clade specific genes across all 21 *Mab* isolates (ignoring genes with low-confidence calls, marked with asterisk). The phylogeny of the 21 isolates is depicted above with the *massiliense* clade (includes all *massiliense* isolates) in yellow, the *abscessus* clade 1 in blue and *abscessus* clade in green. The genes highlighted in gray are genes seen in lineage specific gene essentiality of Figure 5 in the main text.

### Confidence Assessment of Essentiality Calls by HMM

One difficulty in assessing the certainties of the HMM calls is that while the formal state probabilities are calculated at individual TA sites, the essentiality calls at the gene level are made by taking a majority vote (the most frequent state among its TA sites). This does not lend itself to formal certainty quantification.

In the output files of the HMM, we include the local saturation and mean insertion count for each gene, along with the gene-level call and state distribution. Short genes have been observed to be more susceptible to being influenced by the essentiality of an adjacent region, which is evident by examining the insertion statistics. For example, consider a hypothetical gene with just 2 TA sites that is labeled as ES by the HMM, but has insertions at both sites. It might be explained by proximity to a large essential gene or region, due to the "smoothing" the HMM does across the sequence of TA sites. Thus, we can sometimes recognize inaccurate calls by the HMM if the insertion statistics of a gene are not consistent with the call (i.e. a gene labeled as NE that has no insertions, or conversely, a gene labeled ES that has many insertion).

In our paper on the HMM in Transit [4], we showed a plot of random samples from the joint posterior distribution of local saturation and mean insertion counts (at non-zero sites) for the 4 essentiality states, which nicely demonstrates that ES genes have near-0 saturation and low counts at non-zero sites, NE genes have high saturation and counts, GD genes fall in between, and GA genes are almost fully saturated with excessively high counts.

Following this idea, we can use the observed insertion counts in each gene to assess the confidence in each of the essentiality calls by the HMM. Rather than modeling them as 2D distributions, we *combine* them into 1D Gaussian distributions over the *overall mean insertion count* in each gene, including sites with zeros (Figure SM4). The mean count for essential (ES) genes usually around 0, typically around 5-10 for growth-defect (GD) genes, around 100 for non-essential (NE) genes, and >300 for growth-advantaged (GA) genes (assuming TTR normalization, by default).

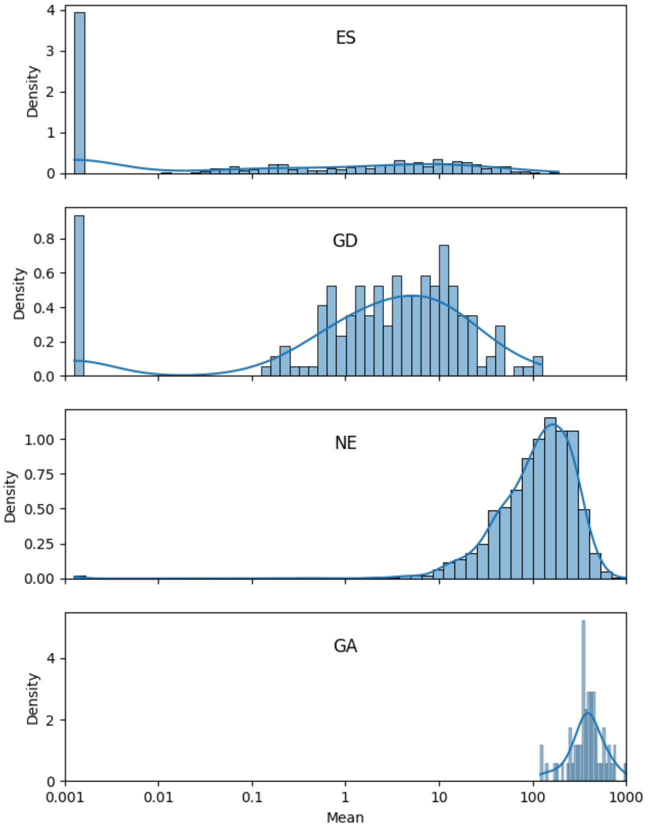


**Figure SM4.** Insertion statistics for genes with different essentiality categories by HMM calls.

The HMM routine in Transit was modified to calculate these conditional distributions empirically for each dataset on which the HMM is run and use it to assess the "confidence" in each of the essentiality calls. First, the mean and standard deviation of saturation is calculated along with insertion count over all the genes in each of the 4 states (ES, GD, NE, and GA). Then, for each gene, the probability density (likelihood) of its mean count is computed with respect to the Normal distribution for each of the 4 states. For example, suppose a gene g is called state s.

$$P(g|s) \ = \ N(cnt(g)|\mu_{cnt}(s), \sigma_{cnt}(s))$$

The 4 probabilities are then normalized to sum up to 1. Finally, the "confidence" of the HMM call for a gene is taken to be the normalized probability of the called state.

This confidence score nicely identifies genes of low confidence, where the local saturation and mean insertion count seem inconsistent with the HMM call. The low-confidence genes are biased toward short genes (with 1-3 TA sites), though they include some large genes with many TA sites as well. Some of the former are cases where the call of a short gene is influenced by an adjacent region. Some of the latter include ambiguous cases like multi-domain proteins, where one domain is essential and the other is not. We observed that there are often "borderline" or ambiguous genes where the called state has significant probability (>0.2), but is not the most probable state (i.e. another state is more likely, based on the insertions in the gene) (see Figure SM5).

Criteria (for gene g with called state c):

- genes where the called state has the highest conditional probability (most likely, given their mean count) are **'confident'**
- genes where $P(g|c) \ > 0.2$, but there is another state that has higher probability are **'ambiguous'**

- genes with $P(g|c) < 0.2$ are **'low-confidence'**

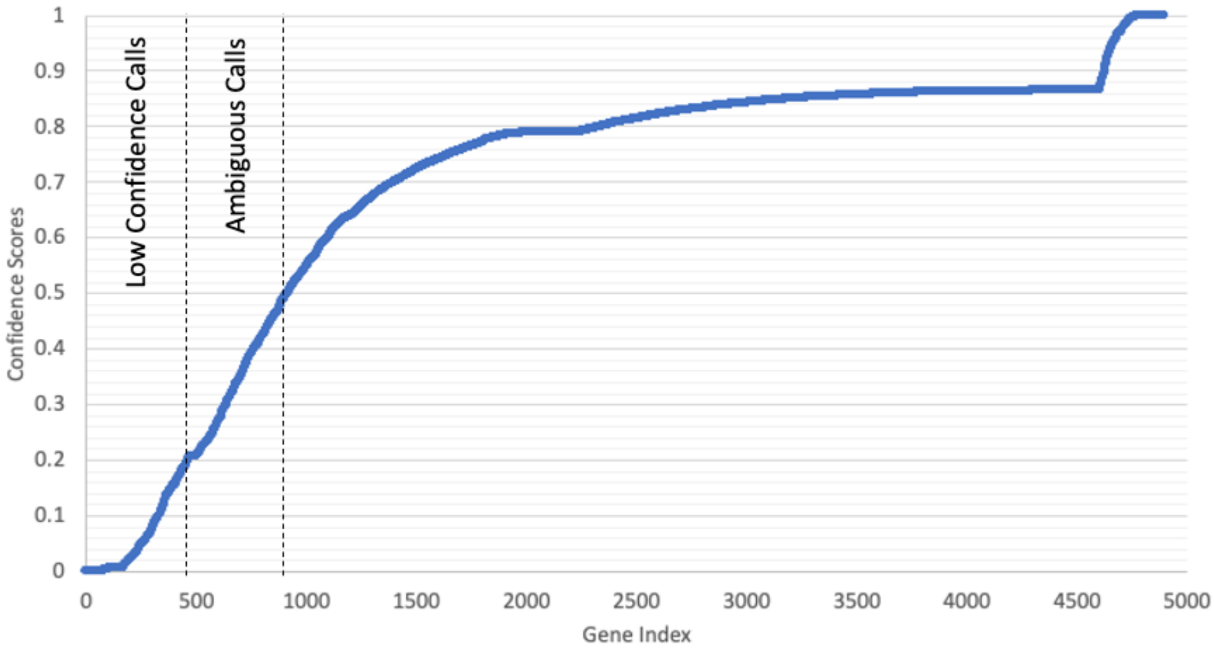


**Figure SM5: Confidence scores of genes in ATCC 19977 strain**. When the 4900 genes in the reference strain are sorted in order of confidence score, most of the genes score near 0.8. The area below the first dashed lines (i.e., confidence score < 0.20) is where genes tend to have a low-confidence score. The area between the two dashed lines is where most of the ambiguous genes reside.

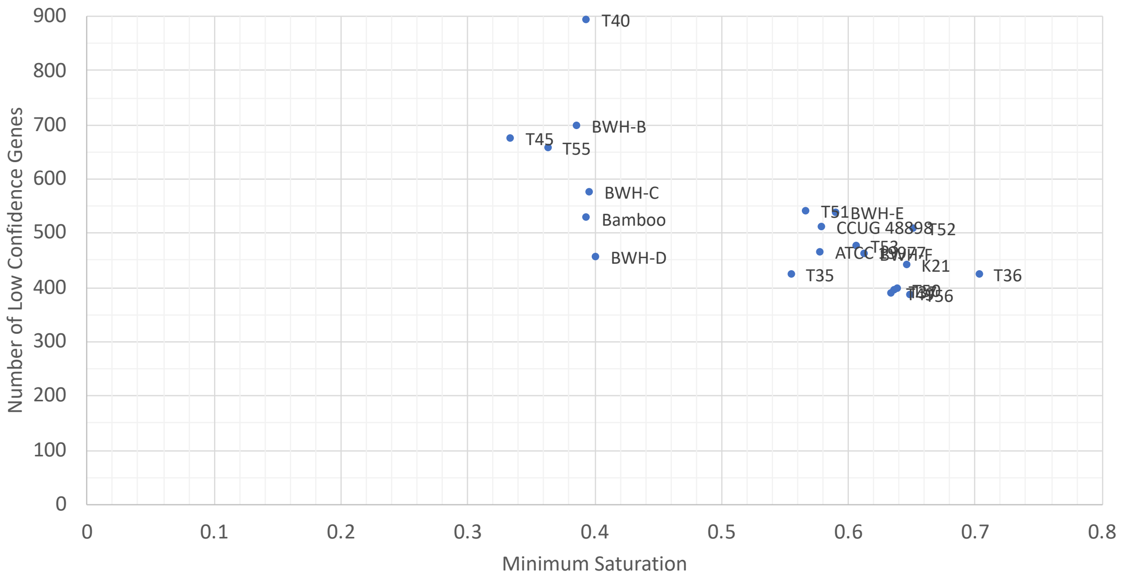


**Figure SM6. Comparison of Saturation of Datasets to Number of Low Confidence Genes.**

Confidence scores can be used to assess the impact of low-saturation datasets on the HMM output. Figure SM6 shows the number of low-confidence genes for each isolate, along with the minimum saturation of the input datasets. The isolates with the lowest saturation have more genes with low confidence scores. Figure SM6 suggests that the minimum saturation replicate for T45 should be dropped, to reduce the noise during the HMM calls. However, when the HMM calls generated using both replicates of T45 are compared to the calls generated using just the high saturation replicate, there is only a minor difference. In fact, 116 more genes have matching essentiality calls to T51 (a close relative on the phylogenetic tree) when using both replicates of T45, then when just using the high saturation replicate (4322 matches vs. 4206 matches). Thus, the addition of a low saturation isolate provides some additional information not present in the high saturation replicate that makes including it in the HMM analysis beneficial.

HMM merges replicates by calculating the average of insertions at each TA site across the replicates of a given isolate. With multiple replicates, even if one replicate has low saturation, the sites with insertions in the merged dataset is the union over the replicates, therefore the saturation is equal to or greater than the maximum saturation of the individual replicates. As seen in Table SM1, the merged saturation of the datsets used in this analysis are 40% or greater.

**Table SM1. Summary of TnSeq datasets, showing the merged saturation* among replicates, as processed by the HMM.**

| Isolate | * Merged Saturation | Minimum Saturation | Maximum Saturation | Average Saturation | Number of Replicates |
|---|---|---|---|---|---|
| CCUG 48898 | 0.580 | 0.580 | 0.580 | 0.580 | 1 |
| ATCC 19977 | 0.750 | 0.579 | 0.641 | 0.614 | 3 |
| BWH-D | 0.683 | 0.402 | 0.628 | 0.515 | 2 |
| T53 | 0.747 | 0.607 | 0.658 | 0.625 | 3 |
| T55 | 0.660 | 0.364 | 0.604 | 0.473 | 3 |
| BWH-B | 0.522 | 0.387 | 0.403 | 0.395 | 2 |
| T36 | 0.703 | 0.704 | 0.704 | 0.704 | 1 |
| T52 | 0.654 | 0.652 | 0.652 | 0.652 | 1 |
| BWH-E | 0.734 | 0.591 | 0.630 | 0.613 | 3 |
| BWH-C | 0.703 | 0.397 | 0.632 | 0.505 | 3 |
| T40 | 0.399 | 0.395 | 0.395 | 0.395 | 1 |
| T37 | 0.789 | 0.637 | 0.690 | 0.658 | 3 |
| T45 | 0.596 | 0.335 | 0.563 | 0.449 | 2 |
| BWH-F | 0.754 | 0.614 | 0.696 | 0.642 | 3 |
| T51 | 0.750 | 0.568 | 0.699 | 0.620 | 3 |
| T50 | 0.785 | 0.640 | 0.714 | 0.673 | 3 |
| T49 | 0.784 | 0.635 | 0.726 | 0.688 | 3 |
| T56 | 0.805 | 0.650 | 0.722 | 0.696 | 3 |
| Bamboo | 0.395 | 0.395 | 0.395 | 0.395 | 1 |
| K21 | 0.648 | 0.647 | 0.647 | 0.647 | 1 |
| T35 | 0.730 | 0.556 | 0.607 | 0.579 | 3 |

*Merged saturation is the saturation after counts are averaged across all replicates for each site

### Sensitivity Analysis of Resampling based on Saturation of Datasets

We have empirically found resampling to work well over a range of saturations through many TnSeq studies (spanning 25-75%), with at least some of the hits being expected or making sense biologically (for each experiment), and in some cases being validated. The normalization procedure used (TTR; [5]) automatically adjusts for differences between saturation of individual datasets by inflating the mean counts in less saturated samples to compensate. This preserves the expectation of the mean insertion count in each gene.

To quantify the impact of saturation on resampling, we conducted a simulation experiment using a high-quality dataset consisting of 6 replicates of *Mtb* H37Rv *in-vitro* (grown on 7H10 medium) and 6 replicates *in-vivo* (inoculated into C57BL6 mice) [6]. These datasets had an acceptable saturation of 47-57% (average = 55.2%). Running resampling between all 6 *in-vitro* and 6 *in-vivo samples*, 187 conditionally essential genes were identified, taken to be the gold-standard (or "true" essentials) for the purposes of this experiment. Artificially sparsified versions of these datasets were created by randomly setting a fraction of TA sites to 0. Importantly, sites were zero-ed out independently for the *in-vitro* and *in-vivo* groups, to emulate the study of *Mab* isolates, in which comparisons between isolates necessarily involved comparison between two independent transposon libraries. Since insertions in non-essential regions are stochastic in each library, the missing sites would be expected to be different between strains (but the same between replicates within a strain). Then, 3 *in-vitro* replicates and 3 *in-vivo* replicates were chosen at random and analyzed for conditionally essential genes using resampling. This was repeated 5 times. Finally, the significant number of genes for each level of sparsification were examined and categorized as TP, FP, TN, or FN, compared to the significant genes obtained by resampling all 6-vs-6 replicates at their original saturation.

As can be seen in Figure SM7, the detection of True positives (relative to full saturation 6-vs-6 comparison of *in-vitro* vs *in-vivo*) decreases gradually with decreasing saturation. However, the false positives (non-essential genes according to the gold-standard that are classified as essential) begin to increase substantially below about 38% saturation. The dashed line is the point at which TP surpasses FP, saturation = 37.5%. The error bars show the range over 5 trials.

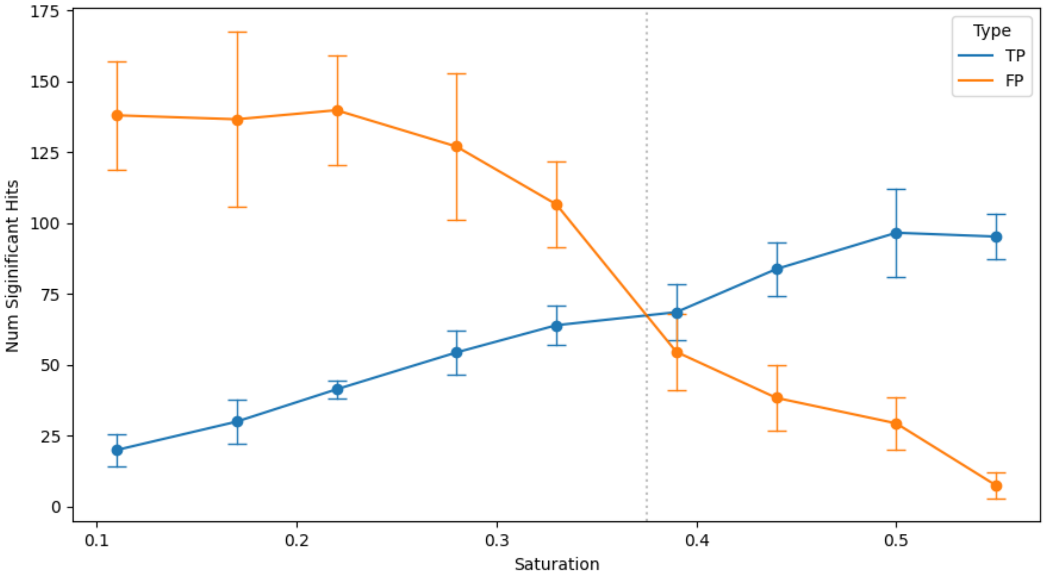


**Figure SM7. False Positives and True Positives at each saturation level.** Three replicates were selected *in-vitro* and three replicates were selected *in-vivo* for each sparsification level. The resampling results are compared to the resampling output of all six replicates of *in-vitro* vs. all six replicates *in-vivo* (with no adjustments to saturation).

In the clinical isolate datasets, two replicates fall below the cut off 38%. One T45 replicate has a saturation of 33.5%. However, T45 has another replicate with 55.3% saturation, which is adequate. The other is for T55; one replicate has a saturation of 36.5%, but the saturation of the other two replicates are 45.0% and 60.4%.

### Sensitivity Analysis of ANOVA to Saturation of Datasets

With ANOVA, the primary risk due to inclusion of low-saturated datasets is increased rate of false positives. To address this, artificially sparsified replicates of the reference strain were used to assess the effect of saturation in the replicates on significantly variable genes detected by ANOVA. The saturations of the reference strain replicates are 0.579, 0.623, 0.641. One, two and all three of the replicates of the reference strains were sparsified by randomly setting a portion of the insertion counts at TA sites (the same sites across all replicates being adjusted) to 0. ANOVA was re-run for all isolates with these sparsified datasets for the reference strain.

As seen in Figure SM8, when there is only one replicate affected by low saturation (< 35%) the combined saturation is still high and mean insertion count in each gene across replicates in the reference sample is not greatly affected. Whereas if all replicates have very low saturation, the mean of genes in the reference sample will appear to decrease, resulting in ANOVA detecting more false positives. Thus, ANOVA is tolerant of individual replicates with low saturation, so long as there are other high saturation replicates for that isolate. It is more likely to generate false positives if the max saturation in the replicates for an isolate is low. As seen in Table SM1, all isolates included in our analyses have a max saturation of 40% or greater, thus our analyzes have less likelihood of reporting false positives.

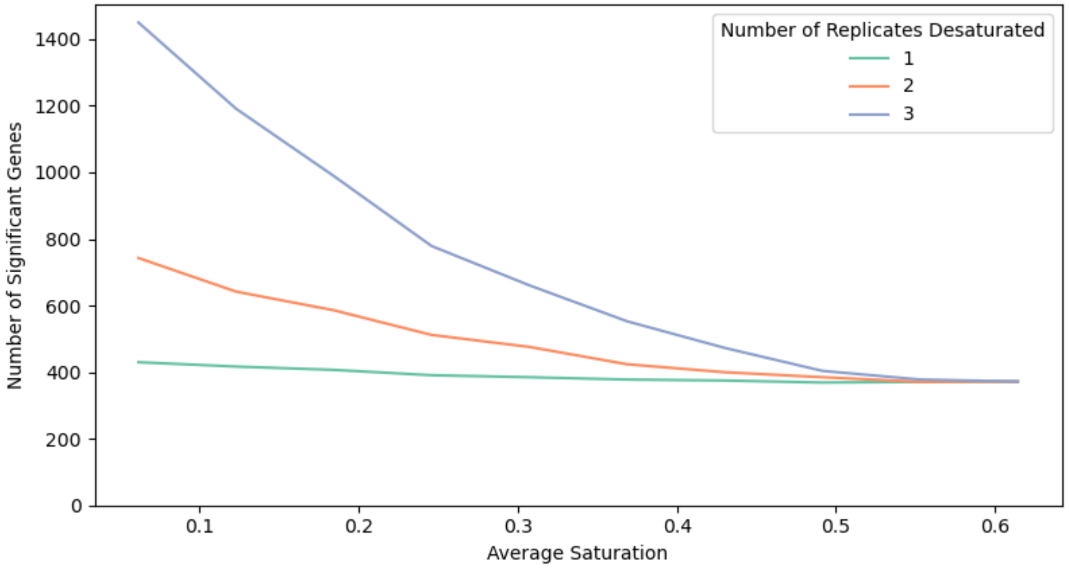


**Figure SM8. The effect of saturation of the reference strain on ANOVA.** We sparsified one, two and three replicates of the reference strain. For each adjustment, ANOVA is run on all 21 isolates, with the adjusted reference library. ANOVA is unaffected by a single low saturation replicate. However, when all three replicates have low saturation, the number of false positives increase.

**Plasmids**

Plasmids have been previously reported in multiple *M. abscessus* isolates, some carrying drug-resistance or toxic metal-tolerance genes [7, 8]. In Dedrick, Aull [9], plasmids were observed in approximately half of the Mab isolates evaluated in their study. To identify plasmids within our collection of Mab clinical isolates, we employed a two-pronged approach. First, we calculated the mean coverage for each contig in the de-novo assembly of each isolate (using Abyss), looking for contigs with inflated mean coverage (≥ 3 standard deviations above the genome-wide weighted mean coverage, weighted by contig lengths) and length ≥ 10 kb. Most mycobacterial plasmids have low-copy number (typically 2-3x relative to the genome-wide mean coverage), and this was also observed by Dedrick, Aull [9]. Nine out of 26 isolates in our collection had contigs with coverage in the 1.7-3.1x range, with sizes 11-104 kb, which is typical for most plasmids (see Supplemental Table 2). As a positive control, we identified a 23 kb contig in the ATCC 19977 sequencing data (with coverage 3.1-fold higher than the rest of the genome), corresponding to the known plasmid pMAB23 that was published for this strain [7].

Secondly, we performed a homology search of the ORFs in each of these high-coverage contigs using Blast against a database of 483 mycobacterial plasmids (downloaded from Genbank) to determine whether they contain genes necessary for replication. We consistently identified orthologs of 5 plasmid-related genes: RepA (binds ori), ParA/Soj (chromosome partitioning protein), a recombinase/resolvase, an Abi-like protein, and MobF (relaxase). 5 of the 9 contigs with excess coverage contained orthologs of all 5 of these genes (ORF ids are listed in Supplemental Table 2), confirming that these are putative plasmids. Mab isolates BWH-B and BWH-E have an 18 kb plasmid; ATCC 19977 has a 23 kb plasmid; Mab-T56 has a 25 kb plasmid; and *M. bolletii* T42 has a 42 kb plasmid. The 4 other contigs with excess coverage contain transposase and/or phage genes, representing other mobile elements.
